# LONGITUDINAL CHANGES IN WHITE MATTER MICROSTRUCTURAL STATUS FOLLOWING QUANTIFIED HEAD-BALL IMPACTS IN SOCCER: A PRELIMINARY, PROSPECTIVE STUDY

**DOI:** 10.1101/2025.02.14.637974

**Authors:** Hugh McCloskey, Carolyn Beth McNabb, Pedro Luque Laguna, Bethany Keenan, John Evans, Derek K Jones, Marco Palombo, Megan Barnes-Wood, Rhosslyn Adams, Sean Connelly, Peter Theobald

**Affiliations:** Cardiff School of Engineering, Cardiff University, The Parade, Cardiff, CF24 3AA, UK; Cardiff University Brain Research Imaging Centre, Cardiff University, Maindy Road, Cardiff, CF24 4HQ, UK; Charles Owen & Co, Croesfoel Industrial Park, Wrexham, LL14 4BJ, UK; Football Association of Wales (FIFA Medical Centre of Excellence), Hensol, Pontyclun, CF72 8JY, UK

## Abstract

Repetitive, sub-concussive head impacts have been associated with increased chronic traumatic encephalopathy (CTE) incidence. CTE diagnosis traditionally relies on post-mortem examination, which limits precise correlation between cause- and-effect. This prospective study embraced innovative diffusion magnetic resonance imaging, which enables in vivo quantification of acute, sub-acute and chronic changes in brain tissue microstructure. This approach was used to evaluate changes in white matter microstructural status at intervals up to 180 days following a specified soccer heading protocol. This study was approved by the university ethics panel. Twelve males (21 – 23 years) were recruited to the study and gave signed, informed consent. Six Intervention participants were university-level soccer players, with 6 Control participants drawn from university-level non-contact sports. Multi-shell diffusion-weighted MRI data were acquired on a 3T Siemens Connectom (300mT/m) scanner using the HARDI protocols. Baseline measures of fractional anisotropy, mean diffusivity and mean kurtosis were acquired at day 0. The Intervention cohort then performed 10 soccer ‘headers’ in a laboratory, with acceleration-time data captured using an instrumented mouthguard and post-processed to report common metrics. The Intervention group was then re-scanned at day 1 (n = 6), day 90 (n = 5) and day 180 (n = 4). The Control group was re-scanned at day 1 (n = 6) and day 180 (n = 3). Many brain tracts were identified as having significant (p < 0.05) changes in white matter microstructural changes at day 90, which correlated strongly with the magnitude of head impact. A smaller number of tracts had changes at day 1 and day 180. These results indicate that, within this pilot population, the magnitude of repeated soccer headers appears to correlate with the magnitude of white matter microstructural change. Additional investigation is required to determine whether the effect of such an intervention influences long-term brain health risk.

## 1. INTRODUCTION

Dementia pugilistica, or ‘chronic traumatic encephalopathy’ (CTE), is a progressive, degenerative brain disease that is typically associated with cognitive and behavioural changes (1). Diagnosis can only be confirmed during post-mortem examination, identified by pathognomonic neurofibrillary tangles and hyperphosphorylated tau protein deposition. Presentation correlates with repeated head impacts, with more severe CTE associated with a longer playing careers of elite male American football players (2), a sport synonymous with helmeted head collisions. CTE has also been identified in some former elite male soccer players, especially those who played in defensive positions, which traditionally demand more head-ball impacts (3)(4). Sport governing bodies now frequently refine protocols and procedures that consider concussion and CTE risk, typically focusing on reducing head impact frequency and magnitude. As an example, the English and Welsh governing bodies for soccer (Football Association; Football Association of Wales) now advise limiting adult players to 10 ‘high force’ headers per week, defined as a ball delivered from further than 35 m (5). The validity of such a threshold in reducing CTE risk is unclear however, given the decades-long lag between impact exposure and post-mortem diagnosis.

Diffusion magnetic resonance imaging (dMRI) offers opportunity for *in vivo* quantification of acute, sub-acute and chronic changes in brain tissue microstructure (6) (7). Innovative dMRI protocols can achieve detection of subtle white matter microstructural changes (8,9). Using these data, diffusion tensor imaging (DTI) and diffusion kurtosis imaging (DKI) allow for derivation of fractional anisotropy (FA), mean diffusivity (MD), and mean kurtosis (MK). Combining DTI and DKI enables a more detailed characterisation of the diffusion-weighted signal by capturing both Gaussian and non-Gaussian diffusion components, providing greater sensitivity to microstructural alterations such as those observed in CTE. Unlike DTI, which primarily relies on lower b-value data and assumes Gaussian diffusion, the addition of DKI allows for a more comprehensive assessment of tissue microstructure (10). DTI studies have shown that, following mild traumatic brain injury (mTBI), FA values tend to decrease, while MD measures generally increase, suggesting a disruption to barriers hindering diffusion of water molecules (11,12). These changes to FA and MD may reflect axonal loss, demyelination, axonal swelling, and oedema (12–14)

DKI can detect more subtle changes in tissue compartments resulting from mTBI, by quantifying the non-Gaussian component of the diffusion displacement profile with a 4^th^ order kurtosis tensor and deriving rotationally-invariant quantitative measures. MK is one such scalar DKI measure that quantifies the degree of non-kurtosis across all directions in a specified voxel (15). Axonal swelling increases the intra-axonal volume, creating a more restricted environment and leading to more non-Gaussian behaviour that elevates MK (13). Alterations in myelin integrity and extracellular geometry introduces diffusion barriers at various scales and increases diffusion heterogeneity, whilst reactive astrogliosis increases the structural complexity and barriers to diffusion. Lower MK coincides with reduced FA and increased MR at the acute and sub-acute phases, after mTBI (16–18).

Imaging protocols provide a valuable route for quantifying white matter microstructural changes; however, surveying specific populations can have scientific, logistical, and economic barriers. Affordable, wearable sensors can now capture head kinematic data, enabling brain trauma predictions via established injury functions (19)(20). Instrumented mouthguards (iMG) capture six-degrees-of-freedom kinematics during sporting activity and have been widely used in sports, including soccer (21,22). Injury metrics including peak angular velocity (PAV), peak linear acceleration (PLA) and peak angular acceleration (PAA) are direct outputs. Post-processing with respect to the time domain produces head injury criterion (HIC) and rotational injury criterion (RIC), more accurate head trauma metrics.

This study sought to investigate the association between the *magnitude* of head injury metrics (as measured via iMG) and the *magnitude* of change in white matter microstructural status (as measured via DKI), across an age- and gender-matched population.

## METHODS

### Study Design and Participants

A controlled cohort study was designed to longitudinally analyse the brain health of soccer players (“Intervention”) exposed to 10 headers within a controlled, laboratory environment. Six Intervention participants were recruited from a university-level soccer team. All were Caucasian, aged 20.1 ± 1 years, height = 1.78. ± 0.06 m, mass = 69.4 ± 7.6 kg. Recruitment targeted those playing positions where head impacts are less likely, to minimise in-season non-recorded exposures. Control participants (n = 6) were race-, age-, and gender-matched, with age = 20.3 ± 1 year; height = 1.79 ± 0.03 m, mass = 76.3 ± 7.9 kg. They were recruited from university teams including tennis and rowing, targeted due to the low head impact prevalence. Participant exclusion criteria included sustaining a concussion during the period of study, having a prior concussion diagnosis, or any contraindication to MRI. The study was approved by the Cardiff University’s Research Ethics Committees in Engineering (2020_ENGIN_PGR_PT-HM_R1) and Psychology (EC.21.03.09.6320RA). All participants provided written, informed consent.

### Patient and Public Involvement

The Football Association of Wales was involved in the design of this methodology. This ensured that the ball delivery distances and heading frequency represented a training scenario. It was also noted that mouthguards are not worn during soccer, and so participants could not be expected to capture data during interim training and match-play sessions. All participants understood the potential influence of any unrecorded head impacts during the study period, agreed with attempts to minimise additional head impacts and to compile an electronic log of any exposures.

### Heading Protocol and Head Kinematics

A laboratory-based intervention methodology was developed to replicate generalised soccer heading scenarios (23). Each Intervention participant headed 3 short (7.65m), 3 medium (15.5m) and 4 long (23m) passes, spaced evenly across a 90-minute test period. Individual iMGs (Protecht; Sports and Wellbeing Analytics, Swansea, UK.) captured head acceleration-time traces over a period of 104 ms (24). The iMG comprised a 3-axis accelerometer sampling at 1 kHz (± 200 g), a 3-axis gyroscope sampling at 0.952 kHz (± 35 rad s^−1^) and an additional accelerometer with a focussed bandwidth (0.5–1 kHz) (24). Linear and rotational velocity, and angular acceleration, were collected in real-time via Bluetooth connection, with post-processing enabling derivation of kinematic metrics including PAV, PLA and PAA. HIC was calculated for a 15 ms time interval, whilst RIC was calculated over a 36 ms window (23).

### MRI Data Acquisition

Multi-shell diffusion-weighted MRI data were acquired on a 3T Siemens Connectom (300mT/m) scanner. All participants were scanned at day 0 (Intervention = October 2021, Control = April 2022). The six Intervention participants then performed 10 headers, before being re-scanned on day 1 (mean time after intervention = 22 ± 3 hours), day 90 ± 7 and day 180 ± 7. The Control group were re-scanned on day 1 and day 180 ± 7.

High angular resolution data were acquired over multiple shells according to a previously reported protocol (8). This took 18 minutes, using a single-shot spin-echo, echo-planar imaging sequence, with the following directions: b = 200 s/mm^2^ (20 directions); b = 500 s/mm^2^ (20 directions); b = 1200 s/mm^2^ (30 directions) and three shells of 61 directions each at b = 2400 s/mm^2^, 4000 s/mm^2^ and 6000 s/mm^2^. A reverse phase encode, posterior to anterior (P>>A) dataset was also acquired for distortion correction comprising two, non-diffusion-weighted images and b = 1200 s/mm^2^ (30 directions). Data acquisition details for all b-values are as follows: TR = 3000 ms; TE = 59 ms; FOV 220 x 200 x 132 mm^3^; matrix size 110 x 110 x 66; voxel size 2 x 2 x 2 mm^3^; in-plane acceleration (GRAPPA) factor of 2. Diffusion gradient duration and separation were 7 ms and 24 ms, respectively.

### Data Processing and Quality Assessment

#### Pre-Processing

Pre-processing rectifies distortion and artifacts within the MRI data. All non-brain data was excluded by creating a mask of the first non-diffusion-weighted image from each phase-encoding direction, anterior to posterior (A>>P) and P>>A, using the FMRIB Software Library’s Brain Extraction tool (FSL, version 6.0.3) (25). Noise level assessment and noise reduction in the diffusion MRI data was performed using MRtrix3 (www.mrtrix.org), based on principal component analysis (MP-PCA) (26–28). Drifts in scanner performance (and therefore image intensity) were corrected by adjusting the diffusion-weighted MRI data to align with temporary distributed b0 s/mm^2^ (non-diffusion-weighted) images, which were dispersed over time, using a tailor-made MATLAB code (MATLAB R2017b; MathWorks Inc., Natick, Massachusetts, USA). The Slicewise OutLIer Detection (SOLID) approach was also applied, adopting 3.5 and 10 thresholds based on a modified Z-score and a variance-focused intensity metric (29). Adjustment of susceptibility-induced off-resonance fields are deduced from the b0 data acquired in both A>>P and P>>A phase-encoding directions. The necessary corrections for these fields, as well as for the distortions caused by Eddy currents and movements of the subject, are effected through Topup (30,31) and Eddy (32) functionalities (FSL, version 6.0.3). Distortions from gradient non-linearity are addressed through another custom-developed MATLAB code. Attenuation of Gibbs ringing artifacts are corrected using local sub-voxel-shifts technique (28,33), before computing fibre orientation distribution function (both in MRtrix3) to leverage a multi-shell, multi-tissue constrained spherical deconvolution method (28)(34).

#### Quality Assurance

A visual inspection was conducted to identify and discard any data with observable artifacts or improper reconstruction. Head motion was estimated between each pair of consecutively acquired volumes and relative to the first volume in the scan, considering translational and rotational motion in three-dimensional space (35). Data were excluded if displacement exceeded 5.0 mm when compared to the first volume, after motion correction. Additionally, the average values of translational and rotational motions were calculated for each subject, with exclusion for those with average motion metrics exceeding three standard deviations.

### Post-processing

The diffusion kurtosis representation was fitted to the shells up to b = 2400 s/mm^2^ using the Dipy Dkifit model, which fits the diffusion kurtosis model and then computes the diffusion tensor metrics (10,36). This process outputs FA, MD and MK maps for each subject, at each timepoint. The pre-processed diffusion data from each participant’s first timepoint were used to generate bundle segmentations, endings segmentations, and tract orientation maps. TractSeg, a convolutional neural network-based method, was employed for this purpose (37). Briefly, fields of fibre orientation distribution function (fODF) peaks were extracted from the pre-processed diffusion MRI data and estimated using the multi-shell, multi-tissue constrained spherical deconvolution (CSD) method, implemented in MRtrix3 (34). Orientation maps describing fibre orientation within 50 segmented tracts were derived from the same fODF data and incorporated into the segmentation outputs. These are detailed in Supplementary Table 1. Bundle-specific tractograms were then generated using the segmented bundles from the initial timepoint and the fibre direction information derived from the orientation maps. Quantitative tractometry analysis was then performed on the white matter tractograms and parametric maps in TractSeg, using a methodology developed in (38). The tractometry analysis reports metric values at 98 segments along each length.

### Statistical Analysis

Linear mixed effects model analyses were performed using R (version 4.0.3, www.r-project.org) LmerTest and lmerPerm 0.1.9 (39). Pearson’s and Spearman’s correlation, and linear regression, were performed using Python (version 3.8.13, www.python.org) with the scipy.stats package (40).

#### Linear Mixed-Effects Modelling

A linear mixed-effects model (LMEM) was fitted to the data using the LmerTest package, enabling direct comparison of variation at different timepoints across the Control and Intervention groups, including the interaction between time points and group memberships, whilst accounting for random intercepts per subject. Ten thousand permutation tests were then performed to estimate p-values for the fixed effects (lmerPerm 0.1.9) (39). P-values for each tract were corrected for multiple comparisons based on the number of tracts using the Bonferroni method, due to the small sample size and risk of type II error.

#### Longitudinally Correlating the Magnitude of Head Impact with Microstructural Change

The mean and median FA, MD and MK across all tracts were calculated, then the difference between two time-points computed to provide a ‘mean delta’ and a ‘median delta’, in addition to the peak and mean kinematic metrics, for each participant. Potential relationships were analysed through visualisation using scatter plots (generated by the matplotlib library (41)), linear regression, parametric Pearson’s correlation and non-parametric Spearman’s rank correlation (employing the scipy.stats.linregress, SciPy. Stats pearsonr and SciPy. Stats spearmanr functions (42)). Pearson’s correlation coefficient (*R)*, Spearman’s Rho, associated p values and coefficient of determination (*R*²) were calculated for each linear fit.

## RESULTS

The Control and Intervention groups were both scanned at day 0 and day 1. The Intervention group underwent scans at day 90 and day 180. One dataset was excluded from day 90 due to excess head movement, and two participants did not attend the final scan. Three participants from the Control group were also scanned at day 180. This is all described pictorially in Supplementary Figure 1. None of the participants reported any other head impact activity across the investigated period.

### Fractional Anisotropy (FA)

Raw FA results from the LMEM analysis are presented in Supplementary Figure 2.

Figure 1 describes the significant results of the LMEM, comparing the mean FA changes in the Intervention group at day 1, across all tracts. Significant differences (p < 0.05) were identified in the right and left cingulum.

**Figure 1:**
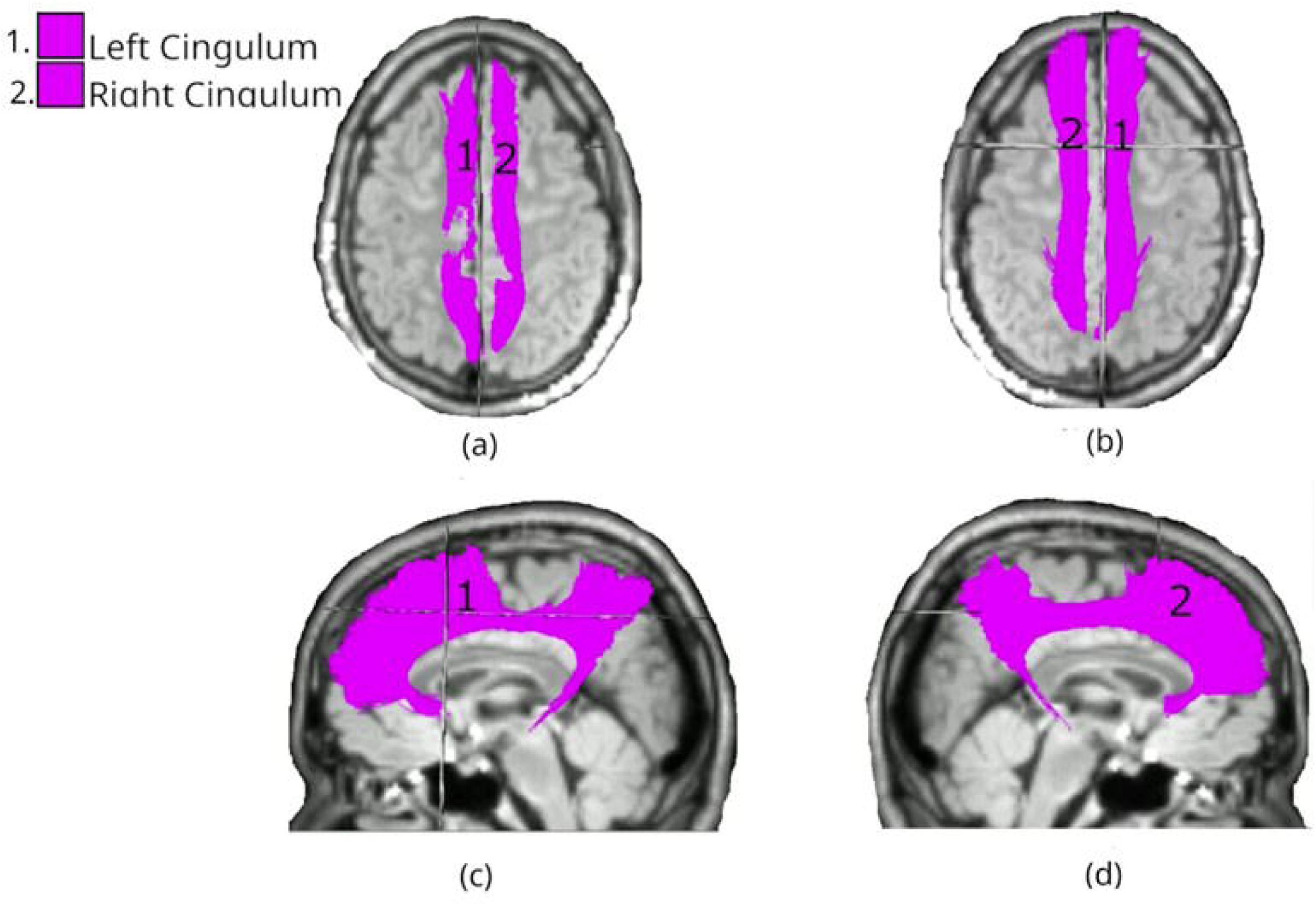
White matter tracts showing significant differences in FA changes between Intervention and Control groups at day 1. Coloured regions overlaid on T1-weighted MRls indicate tracts with statistically significant differences (p < 0.05), based on LMEM. (a) superior axial view; (b) inferior axial view; (c) left midsagittal view; (d) right midsagittal view.

The FA was significantly lower in multiple white matter tracts at day 90 between the Intervention and Control groups (Figure 2). Bilateral reductions were observed in the uncinate fasciculus (UF; left: p=0.010, right: p<0.01), and inferior fronto-occipital fasciculus (IFO; both left and right: p<0.01). Considering the commissural tracts that cross hemispheres, significant differences were found in multiple segments of the corpus callosum (CC), specifically CC_3 (p=0.036), CC_4 (p<0.01), CC_5 (p<0.01), and CC_7 (p<0.01). The right hemisphere showed changes in the anterior thalamic radiation (ATR; p<0.01), inferior longitudinal fasciculus (ILF; p<0.01), and SLF_III (p=0.029). The spatial distribution of these changes suggests a particularly strong involvement of association fibres, especially those connecting frontal and temporal regions, along with substantial involvement of interhemispheric connections through the CC and cerebellar pathways.

**Figure 2:**
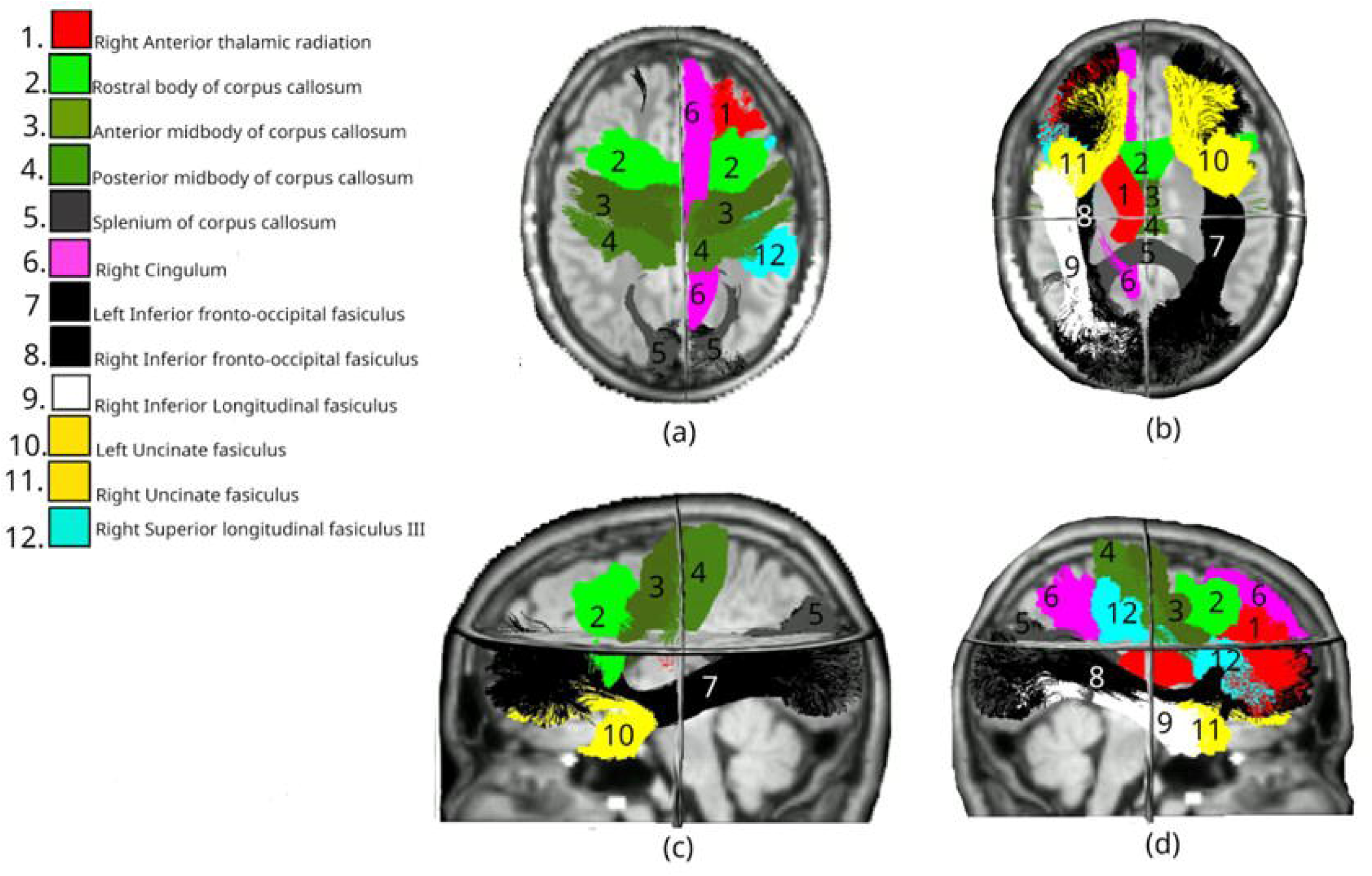
White matter tracts showing significant differences in FA changes between Intervention (day Oto day 90) and Control (day Oto day 180) groups. Coloured regions overlaid on T1-weighted MRls indicate tracts with statistically significant differences (p < 0.05), based on LMEM. (a) superior axial view; (b) inferior axial view; (c) left midsagittal view; (d) right midsagittal view.

All changes identified in the Intervention group had resolved by day 180, with no significant differences between groups in any of the tracts.

### Mean Diffusivity (MD)

Raw MD results from the LMEM analysis are presented in Supplementary Figure 3.

Day 1 MD data revealed significant differences in more than one-third of all tracts (Figure 3). Bilateral differences were identified in the superior longitudinal fasciculus across all three segments (SLF_I, SLF_II, and SLF_III; all p<0.01), cingulum (CG; both left and right: p<0.01), and arcuate fasciculus (AF; left: p<0.01, right: p=0.018). Temporal projections showed bilateral involvement with significant differences in the temporal premotor tract (T_PREM; left: p=0.032, right: p=0.013). Among unilateral tracts, changes were observed in the right anterior thalamic radiation (ATR; p<0.01). The left hemisphere showed changes in the inferior longitudinal fasciculus (ILF; p<0.01), superior temporal and premotor tract (ST_PREM; p<0.01), and stratum (STR; p=0.029). The spatial distribution of these changes again suggests widespread involvement of association fibres.

**Figure 3:**
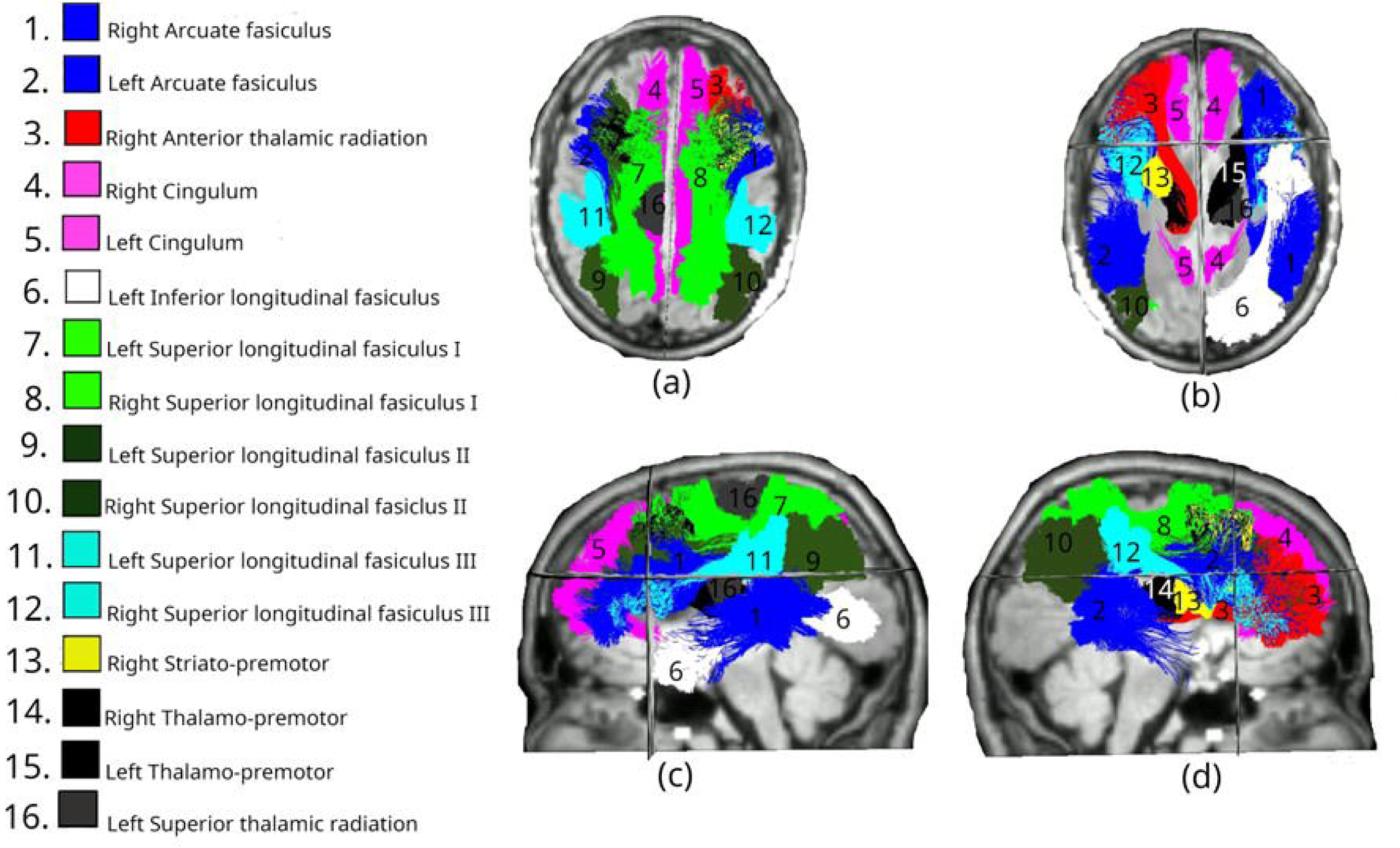
White matter tracts showing significant differences in MD changes between Intervention and Control groups at day 1. Coloured regions overlaid on T1-weighted MRls indicate tracts with statistically significant differences (p < 0.05), based on LMEM. (a) superior axial view; (b) inferior axial view; (c) left midsagittal view; (d) right midsagittal view.

Day 90 data revealed significant differences across 17 tracts, maintaining the bilateral involvement of the SLF_I and CG systems and anterior commissural fibres observed in day 1, though becoming more asymmetric at day 90 (Figure 4). Bilateral changes were observed in the superior longitudinal fasciculus I (SLF_I; both p<0.01), cingulum (CG; both p<0.01) and anterior temporal projections, with both temporal premotor tract and superior temporal premotor tract being affected (T_PREM and ST_PREM left: both p<0.01). Significant differences were found in anterior segments of the corpus callosum (CC_1: p=0.011, CC_2: p=0.008). The right hemisphere showed changes in the anterior thalamic radiation (ATR; p<0.01), inferior longitudinal fasciculus (ILF; p<0.01), uncinate fasciculus (UF; p<0.01), and SLF_II (p=0.033). The left hemisphere showed changes in the arcuate fasciculus (AF; p<0.01), SLF_II (p<0.01), SLF_III (p<0.01), and temporal occipital tract (T_OCC; p=0.027).

**Figure 4:**
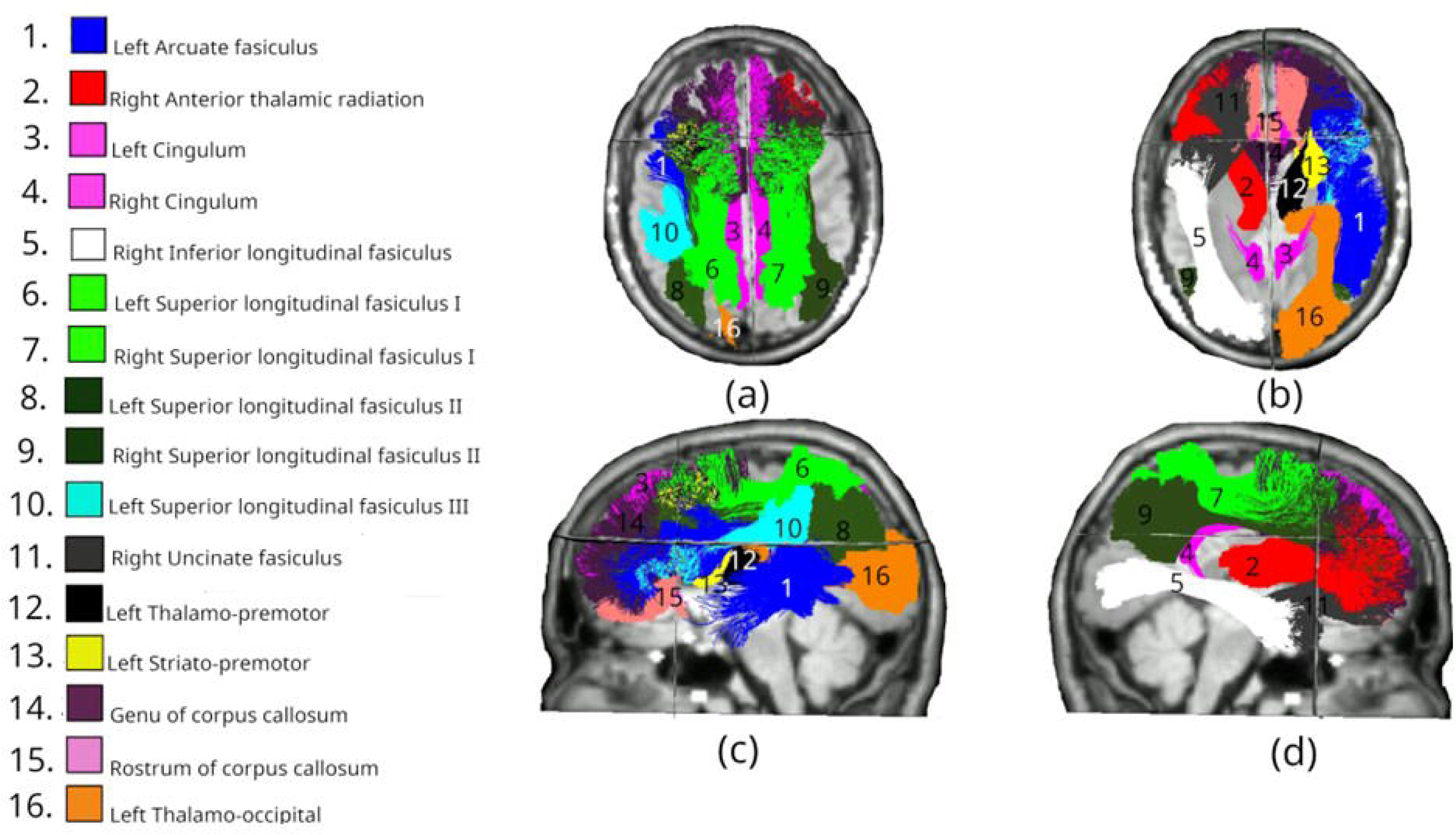
White matter tracts showing significant differences in MD changes between Intervention (day Oto day 90) and Control (day Oto day 180) groups. Coloured regions overlaid on T1-weighted MRls indicate tracts with statistically significant differences (p < 0.05), based on LMEM. (a) superior axial view; (b) inferior axial view; (c) left midsagittal view; (d) right midsagittal view.

Eighteen tracts show a significant MD difference at day 180, the greatest number of the measured period (Figure 5). These showed both persistence and evolution of patterns seen at earlier timepoints. Bilateral changes remained in the superior longitudinal fasciculus I (SLF_I; both p<0.01), cingulum (CG; both p<0.01), superior temporal frontal-occipital tract (ST_FO; both p<0.01), and temporal parietal tract (T_PAR; both p<0.01). The right hemisphere showed changes in the uncinate fasciculus (UF; p=0.026) and stratum (STR; p=0.037). The left hemisphere showed extensive changes in the arcuate fasciculus (AF; p<0.01), inferior longitudinal fasciculus (ILF; p<0.01), SLF_II (p<0.01), SLF_III (p<0.01), temporal premotor tract (T_PREM; p<0.01), stratum (STR; p<0.01), and superior temporal premotor tract (ST_PREM; p=0.014). Compared to earlier timepoints, this pattern maintains the bilateral involvement of SLF_I and CG systems seen at both day 1 and day 90, though shows a marked shift toward left hemisphere dominance in the SLF system. The anterior commissural changes seen at day 90 are no longer present, suggesting a dynamic reorganization of white matter over time

**Figure 5:**
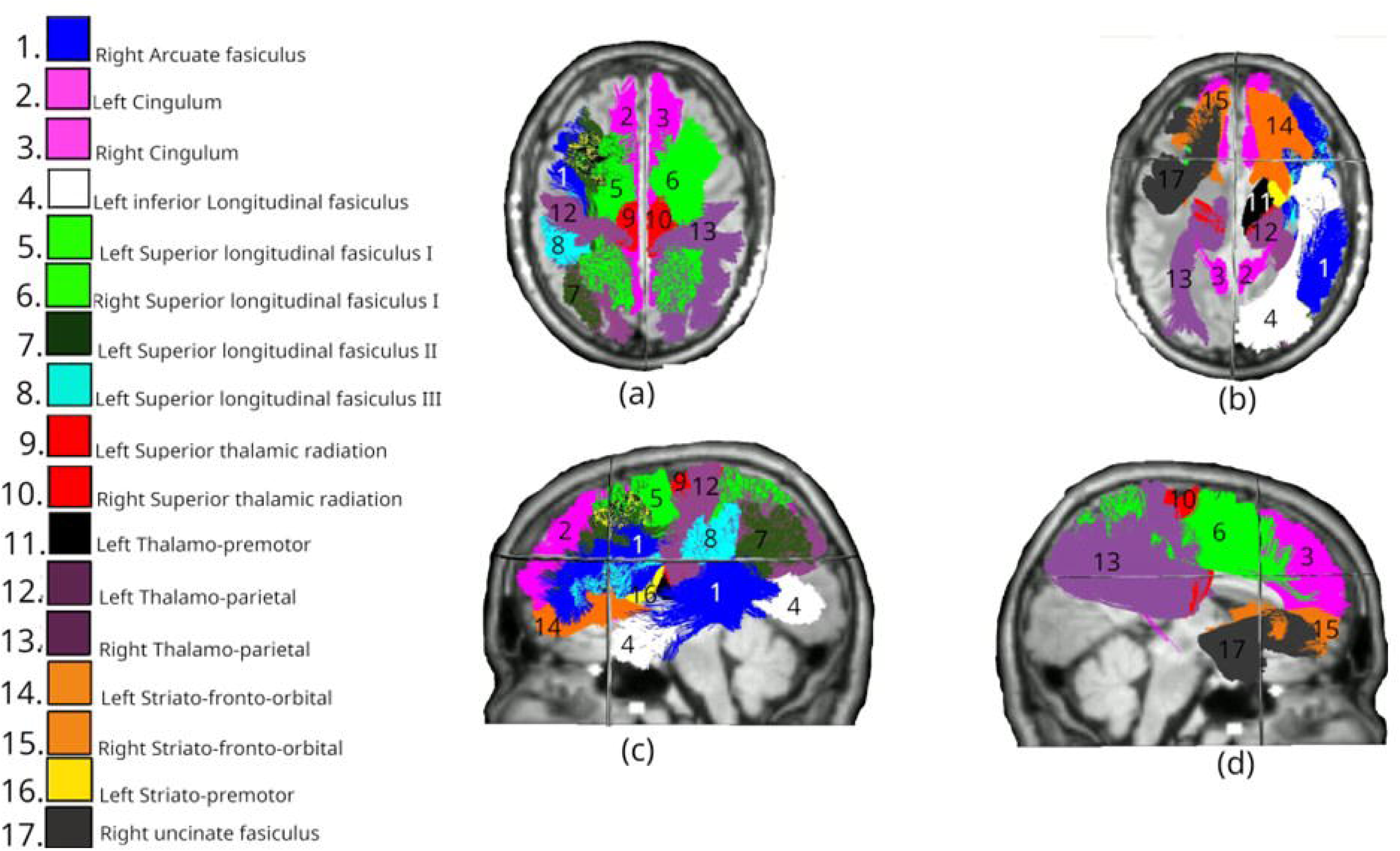
White matter tracts showing significant differences in MD changes between Intervention and Control groups at day 180. Coloured regions overlaid on T1-weighted MRls indicate tracts with statistically significant differences (p < 0.05), based on LMEM. (a) superior axial view; (b) inferior axial view; (c) left midsagittal view; (d) right midsagittal view.

### Mean Kurtosis (MK)

Raw MK results from the LMEM analysis are presented in Supplementary Figure 4.

Analysis of day 1 data revealed a highly focused pattern of significant differences, with changes observed only bilaterally in the cingulum (CG; both left and right: p<0.01). This is the same pattern as day 1 FA (Figure 1).

By day 90 (Figure 6), 17 tracts had significant MK differences. Bilateral changes were observed in the inferior fronto-occipital fasciculus (IFO; both p<0.01) and inferior longitudinal fasciculus (ILF; both p<0.01). Extensive involvement of the CC was also noted, including in CC_2 (p<0.01), CC_3 (p<0.01), CC_4 (p<0.01), CC_5 (p<0.01), and CC_6 (p<0.01). For unilateral tracts, no right hemisphere tracts had a significant difference. Conversely, many left hemisphere tracts showed changes including the uncinate fasciculus (UF; p<0.01), stratum (STR; p<0.01), temporal premotor tract (T_PREM; p<0.01), temporal parietal tract (T_PAR; p=0.006), superior temporal premotor tract (ST_PREM; p=0.008), optic radiation (OR; p=0.034), superior longitudinal fasiculus (SLF_III; p=0.034) and cingulum (CG_left; p<0.01). This pattern shows interesting convergence with both FA and MD findings at day 90, particularly in the involvement of the IFO and commissural fibres, though MK demonstrates unique sensitivity to changes in visual and temporal projection pathways. The expansion from the focused day 1 pattern suggests that while early changes in tissue complexity are limited to the limbic system, by day 90 there is extensive reorganization of tissue microstructure across multiple networks, particularly in the left hemisphere.

**Figure 6:**
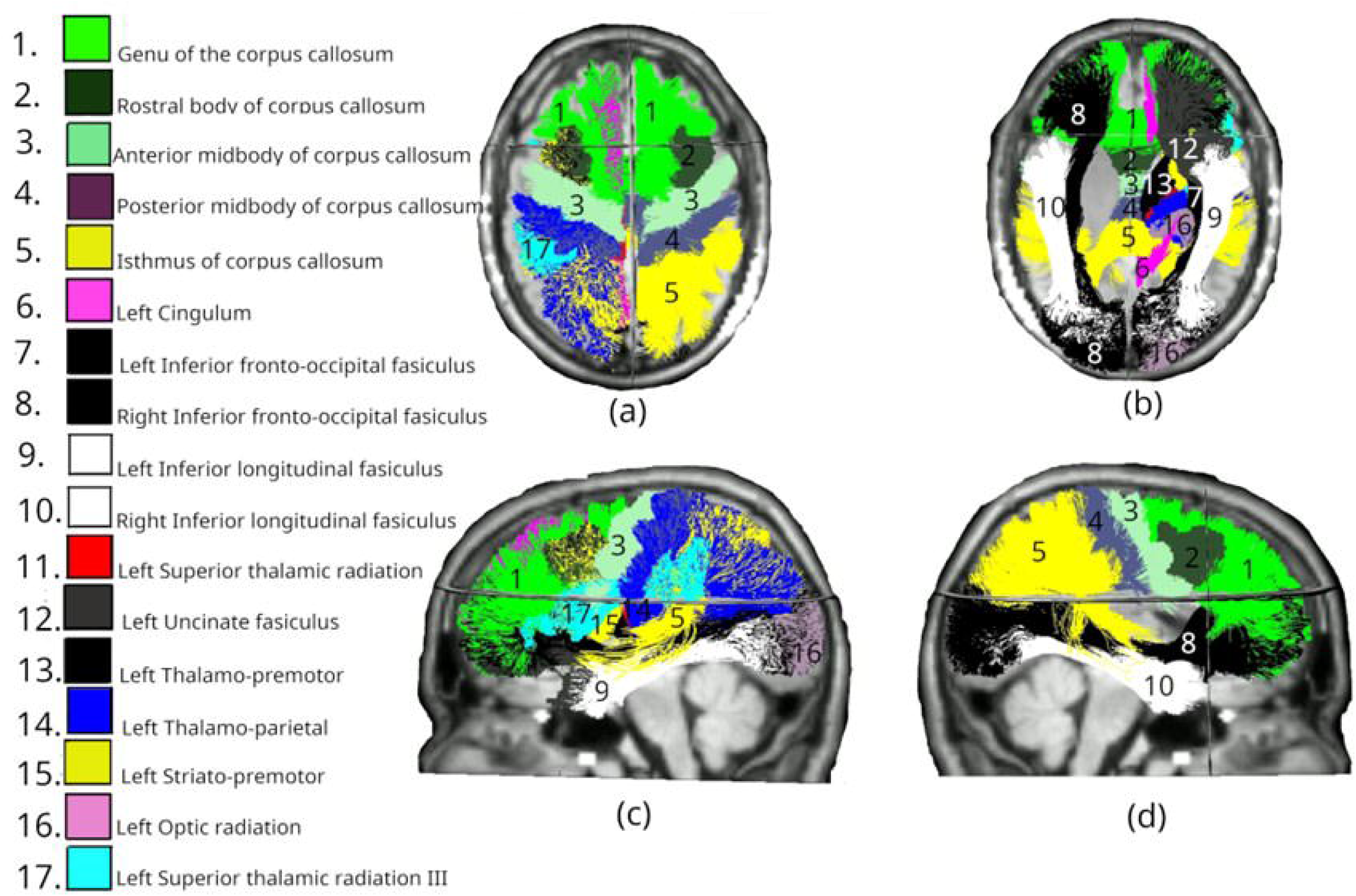
White matter tracts showing significant differences in MK changes between Intervention (day Oto day 90) and Control (day Oto day 180) groups. Coloured regions overlaid on T1-weighted MRls indicate tracts with statistically significant differences (p < 0.05), based on LMEM. (a) superior axial view; (b) inferior axial view; (c) left midsagittal view; (d) right midsagittal view.

Day 180 MK data identified 8 significant tracts, though with a different spatial pattern from day 1 and day 90 (Figure 7). Among commissural tracts that cross hemispheres, involvement of the CC remained prominent but shifted anteriorly, with significant differences in CC_1 (p=0.035), CC_2 (p<0.01), CC_3 (p=0.014), and CC_4 (p=0.026). Bilateral changes were limited to superior longitudinal fasciculus (SLF_I; left: p=0.033, right: p<0.01). Unilateral changes were again absent from the right hemisphere, though observed in the left superior temporal frontal-occipital tract (ST_FO; p=0.015). This pattern shows marked deviation from the extensive bilateral changes seen at day 90 in the IFO, ILF, and temporal projection pathways, suggesting a dynamic reorganization of tissue microstructure over time. Compared to MD at the same timepoint - which showed extensive left-lateralized changes in the SLF system and bilateral involvement of temporal projections, the MK changes are more focused on anterior commissural fibres. The evolution from bilateral cingulum involvement at day 1, through widespread changes at day 90, to this more focused pattern at day 180 suggests a complex temporal progression of microstructural reorganization, with different white matter systems showing distinct trajectories of change over time.

**Figure 7:**
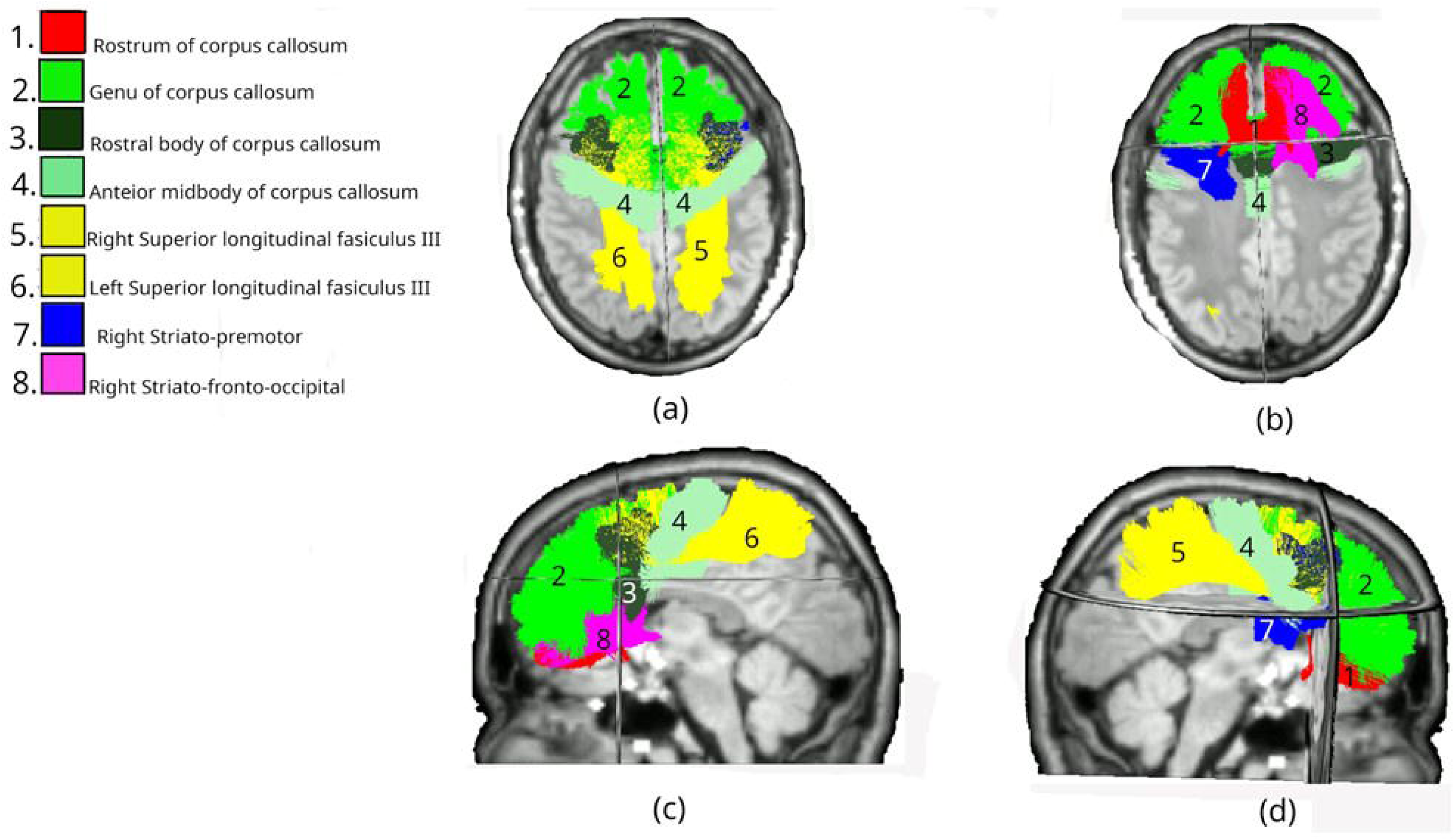
White matter tracts showing significant differences in MD changes between Intervention and Control groups at day 180. Coloured regions overlaid on T1-weighted MRls indicate tracts with statistically significant differences (p < 0.05), based on LMEM. (a) superior axial view; (b) inferior axial view; (c) left midsagittal view; (d) right midsagittal view.

### Kinematic Analysis

Post-processed data describes the peak values, and the cumulative peak values, for each participant (see Supplementary Figure 5). In most instances, the peak and cumulative metrics share common trends: Participant 3 scored highest across all metrics and so was expected to have the highest trauma risk; Participant 4 generally achieved lower metrics and so, assuming all anatomical and physiological variables were broadly comparable across participants, would have a lower injury risk. Participant 2 is notable for their relatively high linear metrics, whilst scoring relatively low rotational metrics.

### Correlating Kinematic injury metrics and DKI metrics

Combining these datasets allows investigating the effect of head impact magnitude on the longitudinal changes in white matter microstructural status, via DKI metrics. DKI metrics are correlated with different kinematic injury metrics at day 1, 90 and 180.

No significant correlations were found between mean and median change in FA and any kinematic metric at day 1. At day 90, a significant negative correlation exists between mean (Fig 8(a)) and median (Fig 8(b)) change in FA, and mean PAV. The significant negative trends indicate that higher mean PAV values during the intervention corresponded with greater reductions in both mean and median FA. Persistent correlations between FA and kinematic metrics exist at day 180, including a strong negative relationship between mean change in FA and mean HIC15 (Supp Fig. 6(a)) and between median change in FA and mean HIC15 (Supp Fig. 6(b)). Additional significant negative correlations were observed between mean and median FA changes, and mean PAV, at day 180 (Supp Figs. 6(c) & (d)).

**Figure 8:**
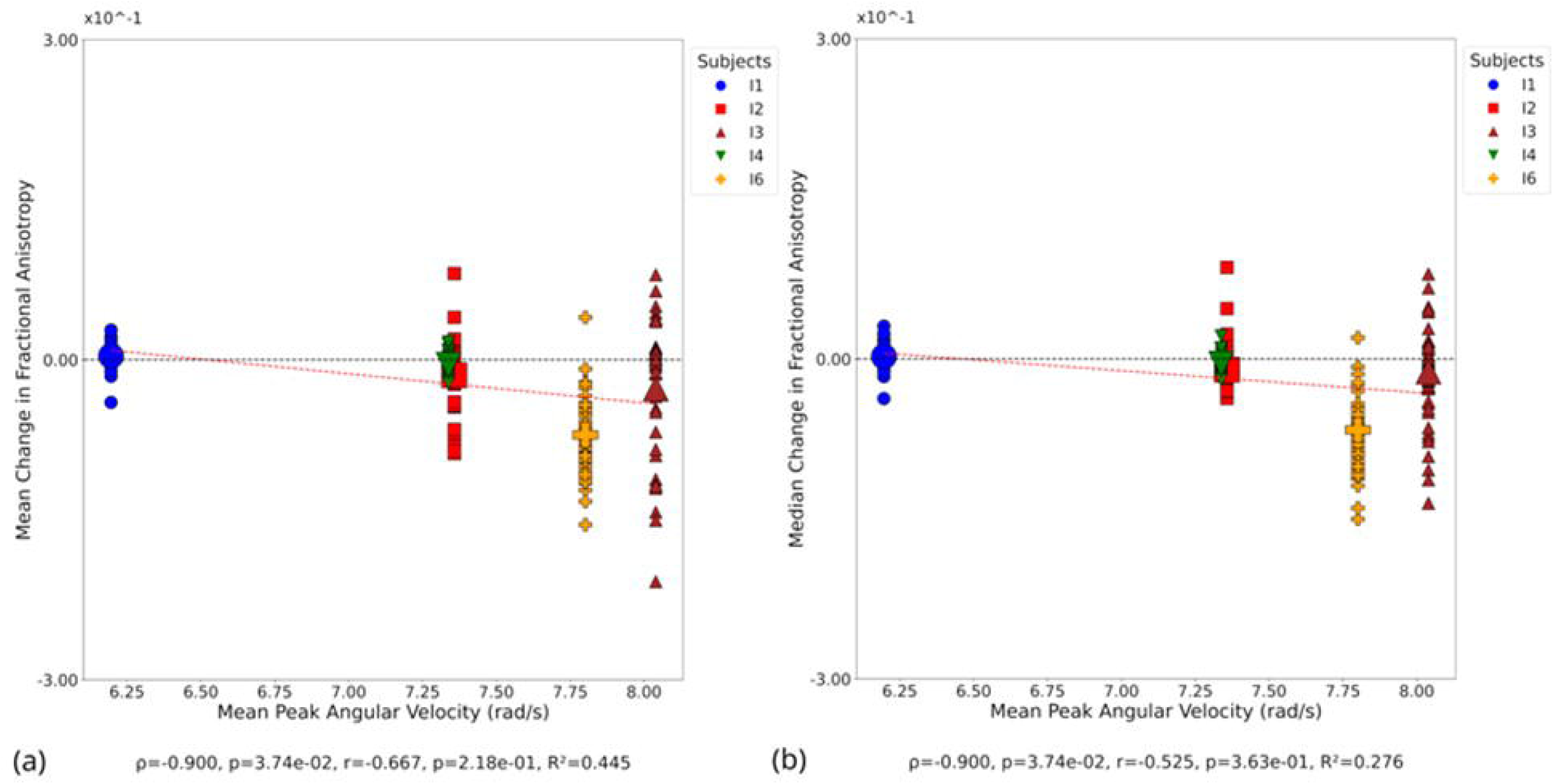
Correlation between FA and kinematic metrics at day 90. Small symbols indicate FA change in individual white matter tracts at the participant level; large symbols indicate overall change in FA for each participant. Statistical significance assessed via Pearson’s correlation (r), Spearman’s correlation (p), and coefficient of determination (R^2^). (a) Mean change in FA versus mean PAV. **(b)** Median change in FA versus mean PAV.

No significant correlation was evident at day 1 between mean change in MD and peak PLA (Supp Fig. 7a), though there was a significant negative correlation between median change in MD and peak PLA (Supp Fig. 7(b)). At day 90, several significant positive correlations were observed between mean (Fig 9(a)) and median (Fig 9(b)) change in MD and peak HIC, mean (Fig 9(c)) and median (Fig 9(d)) change in MD and mean PAV. Additional significant correlations were found between mean (Fig 9(e)) and median (Fig 9(f)) change in MD and mean PLA, and mean (Fig 9(g)) and median (Fig 9(h)) change in MD and peak PLA. At day 180, significant positive correlations were observed between mean change in MD and mean HIC (Supp Fig. 8(a)), and median change in MD and mean HIC (Supp Fig. 8(b)). Strong positive correlations were also found between mean change in MD and peak HIC (Supp Fig. 8(c)), and median change in MD and peak HIC (Supp Fig. 8(d)). Additionally, positive Spearman correlations were observed between mean change in MD and mean PAV ((Supp Fig. 8(e)), and median change in MD and mean PAV (Supp Fig. 8(f)).

**Figure 9:**
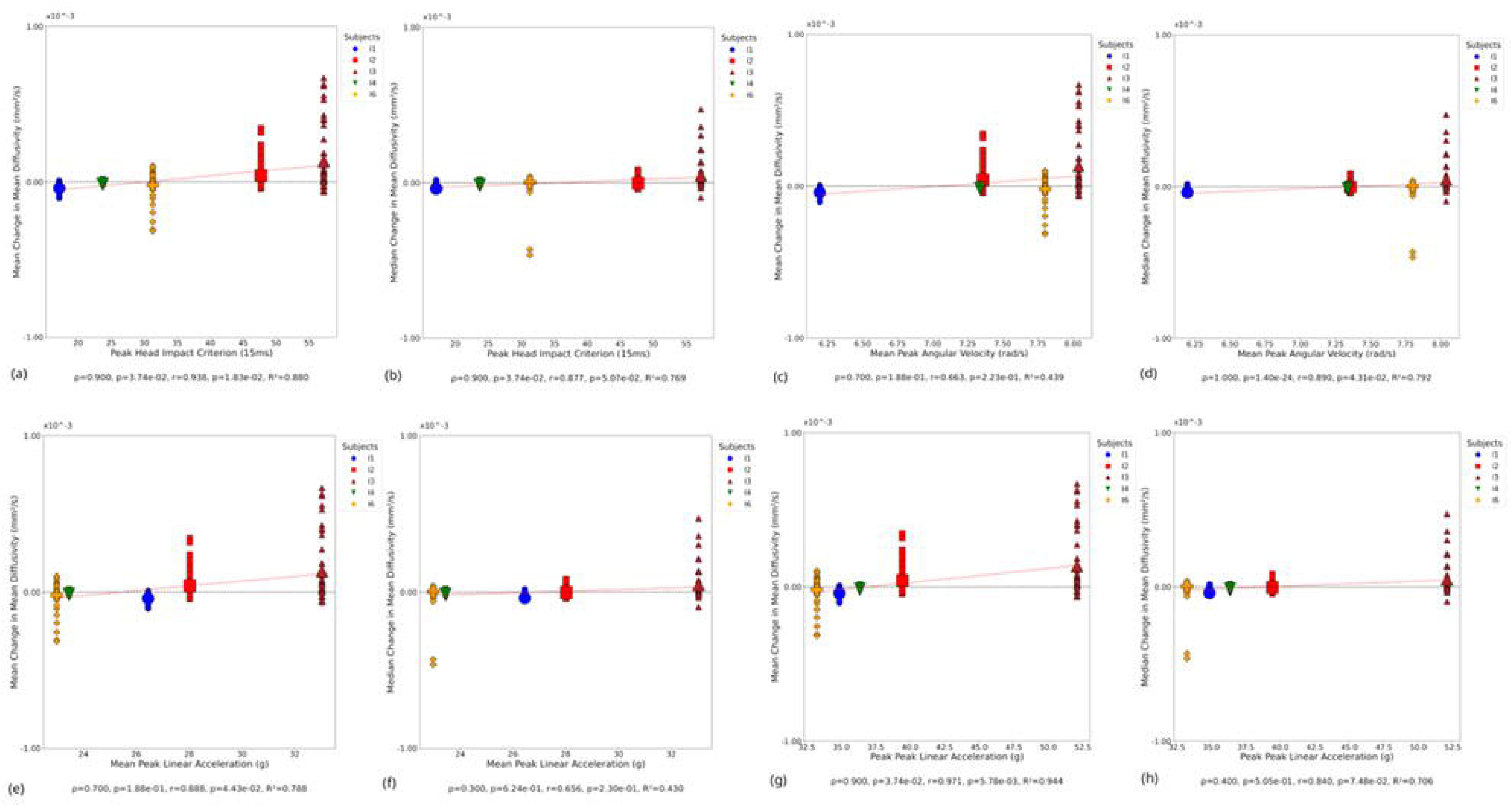
Correlation between MD and kinematic metrics at day 90. Small symbols indicate MD change in individual white matter tracts at the participant level; large symbols indicate overall change in MD for each participant. Statistical significance assessed via Pearson’s correlation (r), Spearman’s correlation (p), and coefficient of determination (R^2^). (a) Mean change in MD versus peak HIC. **(b)** Median change in MD versus peak HIC. (c) Mean change in MD versus mean PAV **(d)** Median change in MD versus mean PAV. **(e)** Mean change in MD versus mean PLA. **(f)** Median change in MD versus mean PLA. **(g)** Mean change in MD versus peak PLA. **(h)** Median change in MD versus peak PLA.

At day 1, significant positive correlations were observed between mean change in MK and mean PAA (Supp Fig 9(a)), while no significant correlation was found for median change in MK and mean PAA (Supp Fig. 9(b)). A significant correlation was observed between mean change in MK and peak PAA (Supp Fig. 9(c)), while median change in MK showed no significant correlation (Supp Fig. 9(d)). At day 90, a significant negative correlation was observed between mean change in MK and mean PAV (Fig. 10(a)), while no significant correlation was found between median change in MK and mean PAV (Fig. 10(b)). Both correlations showed moderate to strong negative relationships based on Spearman’s rho, though only the mean change reached statistical significance, while neither Pearson correlation achieved significance. No significant correlations were observed between MK and any kinematic metric at day 180.

**Figure 10:**
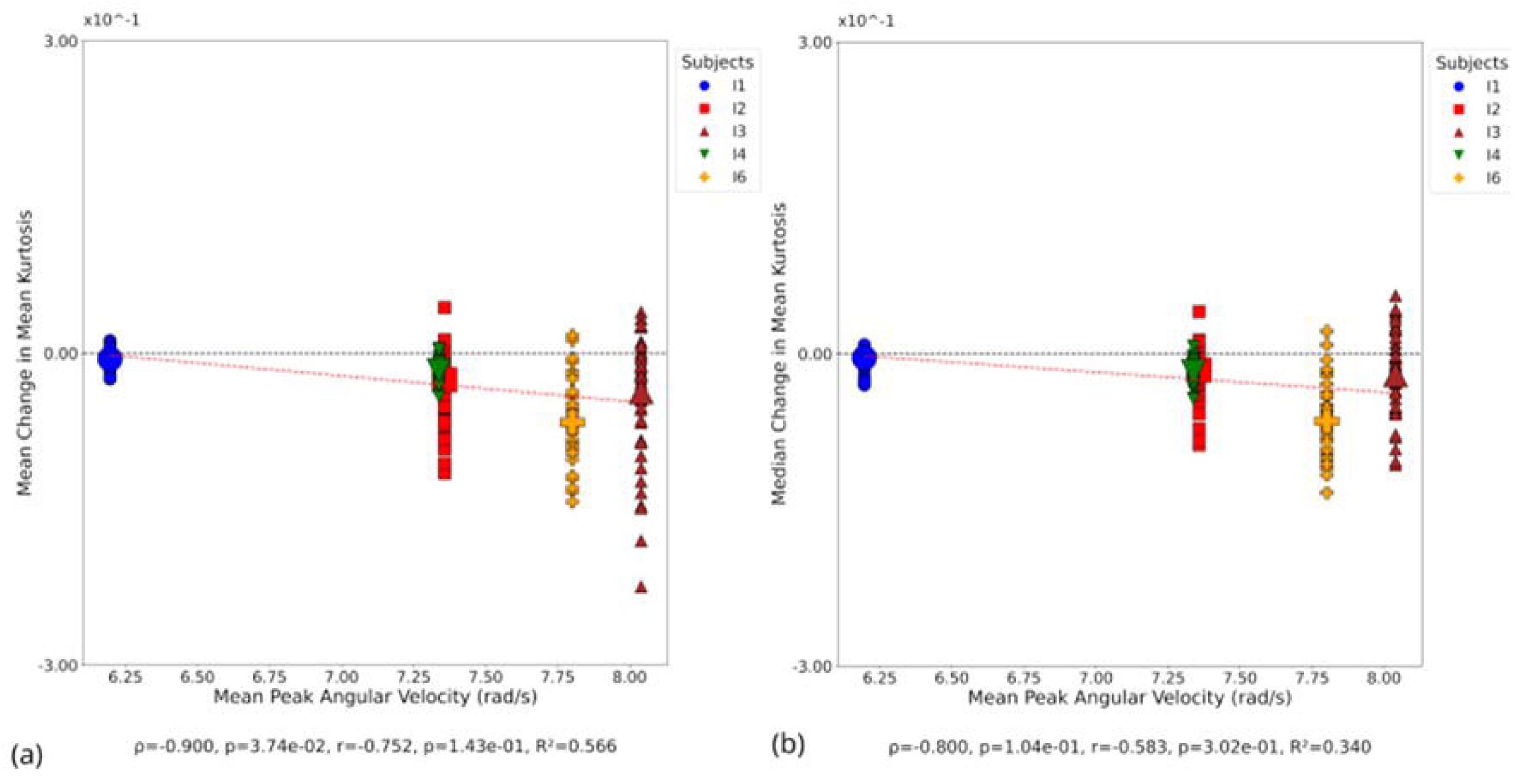
Correlation between MK and kinematic metrics at day 90. Small symbols indicate MK change in individual white matter tracts at the participant level; large symbols indicate overall change in MK for each participant. Statistical significance assessed via Pearson’s correlation (r), Spearman’s correlation (p), and coefficient of determination (R^2^). (a) Mean change in MK versus mean PAV **(b)** Median change in MK versus mean PAV.

## DISCUSSION

This study identified significant differences in changes to DKI metrics in the Intervention versus Control group, tested via LMEM following an intervention of 10 headers, measured across a common period. This study also found statistically significant correlations between the magnitude of kinematic injury metrics and the change in diffusion measures. This was most pronounced at day 90.

LMEM analysis of FA and MK at day 1 demonstrated no statistical change versus the Control group in all tracts except the bilateral cingulum (CG). This lack of change at the acute stage is consistent with previous findings (43). LMEM analysis of MD at day 1 showed significant decreases in the bilateral superior longitudinal fascicle (SLF). A MD decrease in the SLF has previously been observed in concussed athletes (44). Significant decreases were also seen in the anterior thalamic radiation (ATR) and superior thalamic radiation (STR) tracts, a pattern consistent with athletes exposed to collisions during sport (45). The decreasing MD at day 1 coincides with concomitant MD increases in the Control group, which may confound drawing inferences from this change.

The sub-acute (day 90) timepoint LMEM analysis showed statistical decreases in MK and FA, and statistical increases in MD, across many tracts. The spatial location of the changes at this timepoint, with the CC and SLF shown to be preferentially affected, is consistent with patterns of changes associated with mTBI and TBI (45–48)

MK data showing a more marked decrease at day 90, following a slight increase at day 1, is consistent with changes observed in concussed athletes (49). Such changes have been proposed to represent reactive astrogliosis in animal models (50). This MK decrease is also reflective of the trend seen in concussed athletes in the sub-acute phase after injury (18,51).

Intervention FA returned to baseline at day 180 (chronic phase), with no significant differences versus the Control data found in LMEM analysis.

Lower MD and higher MK were identified in many tracts at day 180, coinciding with those that changed at day 90 (sub-acute phase) including the SLF and ILF in MD, and CC and SLF in MK. This biphasic response in MD and MK between the sub-acute and chronic phase in the CC is consistent with a trend observed in other mTBI cases (17).

Tracts of the CC, particularly the most anterior regions, appeared the most significantly influenced by the intervention across the three DKI metrics and across all timepoints, which is consistent with studies reporting concussive and sub-concussive head impacts (52)(17). These patterns in MD and MK have previously been cited as the two most important metrics for detecting repetitive sub-concussive head injuries in athletes (49), and the tracts most likely to be affected were the CC and caudal tract such as the SLF and ILF.

In the currently study HIC15 emerged as the kinematic metric that produced the strongest, and the most, correlations, specifically between peak HIC and MD change at day 90 (R^2^ = 0.880) and day 180 (R^2^ = 0.990). This is consistent with findings from the automotive sector (53). Mean PAV produced the greatest number of significant results across all metrics and timepoints. This is consistent with the PAV reported elsewhere (54). The kinematic metrics presented in this study are likely a conservative representation of soccer exposures, due to all passes <30m, a threshold defining “high energy passes”.

Drawing comparisons between the kinematic values presented in this study and data reported in other studies is challenging, due to variations in measurement protocols, different levels of player training and skill (23). (55) used iMGs to capture head kinematics of university level males (aged 21.3 ± 1.89) and females (aged: 21.25 ± 1.68) with previous soccer experience. The protocol (distance = ∼12.2m, ball velocity = 11.18 ms-1, ball pressure = 0.59 bar) resulted in 78% lower PLA values and 43% lower PAA values, compared to those reported at a similar velocity in this study. Although the protocol setup was similar, the lower values observed are likely due to the reduced ball inflation pressure. (56) used skin patch sensors to capture on-field PLA and PAA values in semi-professional female soccer players. PLA for distances represented in this study (5-20 m) were similar (26.9 g), however PAA values were much higher (5659.8 rad/s2). The use of skin-mounted accelerometers may have contributed to their notably elevated PAA values, as studies indicate frequent occurrence of artefactual data caused by soft tissue deformation. Additionally, it has been observed that females exhibit greater linear and angular acceleration compared to their male counterparts, likely attributable to a lower head mass. Head injury risk metrics varied significantly in youth male footballers (aged 14.8 years), based on ball delivery distance (57). Headers delivered from 15m, simulating in-game scenarios such as a corner kick, resulted in the highest PLA and PAV. Specifically, the attacking headers produced the greatest head impact magnitudes, likely due to the additional power and direction applied to the ball. In contrast, headers from 5m, performed either with the feet grounded or while jumping, produced significantly lower PLA and PAV (58).

## Limitations

The data presented in this study were captured from a very small cohort (n = 12), with analysis and interpretation limited further by participant attrition across the 180-day follow-up. The results are also from a homogeneous population of Caucasian adult males which, whilst intentionally constraining variables, does prevent translation of these findings to consider younger and/or female players. Participant, facility, operator, and financial constraints meant that follow-up dMRI scans were infrequent, though still represent similar intervals to other studies. That Control timepoints were fewer than the Intervention group is consistent too; hence, this study presents a valid and meaningful approach to garner an appreciation of the correlation between head impact magnitude and changes to white matter microstructural status.

## CONCLUSIONS

Statistically significant correlations have been identified between the magnitude of head impacts and the magnitude of change in white matter microstructural status, from day 1 to day 180. Kinematic data were interpreted to represent a relatively modest head impact, though these findings indicate that even these can cause brain changes extending for many months. Aside from sampling larger and more diverse populations, a further step is to determine how cumulative head impacts map on to these correlations, which could provide a greater understanding on the neuropathology underpinning CTE.

## TRANSPARENCY, RIGOUR AND REPRODUCIBILITY

Twelve participants were recruited to this study. All participants were MRI scanned on day 0, and day 1. The 6 Intervention participants were exposed to 10 headers on day 0. Intervention participants were then invited for MRI scanning at day 90, though participant I5 did not attend. Intervention participants were also invited to attend MRI scanning at day 180, though participants I3 and I4 did not attend. The Control participants were invited to attend MRI scanning at day 180, though 3 (C7, C9, C12) did not attend. Participant head kinematics were recorded during the intervention using personalised, instrumented mouthguards. All denture impressions were performed by the same dental surgeon, and all mouthguards manufactured by the same company. All kinematic data was processed using standard methods, to compute established head injury metrics. All MRI data was processed using a linear mixed effects model. Data from this study are available in the Cardiff University repository, with a hyperlink to follow and under a creative commons attribution license (CC-BY 4.0).

## AUTHORSHIP CONTRIBUTION STATEMENT

**Hugh McCloskey:** Validation, Formal analysis, Investigation, Data curation, Writing original draft, Visualisation. **Carolyn Beth McNabb**: Validation, Methodology, Formal analysis, Data curation, Writing – review and editing. **Pedro Luque Laguna**: Software, Formal analysis, Resources, Writing – review and editing. **Bethany Keenan:** Investigation, Data curation, Writing – review and editing. **John Evans:** Investigation, Data curation, Writing – review and editing. **Derek K Jones:** Conceptualisation, Methodology, Writing – review and editing. **Marco Palombo:** Software, Formal analysis, Resources, Writing – review and editing. **Megan Barnes-Wood:** Investigation, Data curation, Writing – review and editing. **Rhosslyn Adams:** Formal analysis. **Sean Connelly:** Investigation. **Peter Theobald:** Conceptualisation, Methodology, Writing – original draft, Writing – review and editing, Supervision, Project administration, Funding acquisition.

## REFERENCES

1. McKee AC, Abdolmohammadi B, Stein TD. The neuropathology of chronic traumatic encephalopathy. Handb Clin Neurol. 2018;158:297–307.

2. Alosco ML, Cherry JD, Huber BR, Tripodis Y, Baucom Z, Kowall NW, et al. Characterizing tau deposition in chronic traumatic encephalopathy (CTE): utility of the McKee CTE staging scheme. Acta Neuropathol. 2020 Oct;140(4):495–512.

3. Mackay DF, Russell ER, Stewart K, MacLean JA, Pell JP, Stewart W. Neurodegenerative Disease Mortality among Former Professional Soccer Players. N Engl J Med. 2019 Nov 7;381(19):1801–8.

4. Russell ER, Stewart K, Mackay DF, MacLean J, Pell JP, Stewart W. Football’s InfluencE on Lifelong health and Dementia risk (FIELD): protocol for a retrospective cohort study of former professional footballers. BMJ open. 2019;9(5):e028654.

5. Premier League. Professional Football Heading in Training Guidance. 2021 Jul 16 [cited 2024 Sep 9]; Available from: https://www.thefa.com/-/media/thefacom-new/files/rules-and-regulations/2021-22/heading-guidance/professional-football-heading-in-training-guidance-summary---july-2021.ashx

6. Aoki Y, Inokuchi R, Gunshin M, Yahagi N, Suwa H. Diffusion tensor imaging studies of mild traumatic brain injury: a meta-analysis. J Neurol Neurosurg Psychiatr. 2012 Sep;83(9):870–6.

7. Toth A, Kovacs N, Perlaki G, Orsi G, Aradi M, Komaromy H, et al. Multi-modal magnetic resonance imaging in the acute and sub-acute phase of mild traumatic brain injury: can we see the difference? J Neurotrauma. 2013 Jan 1;30(1):2–10.

8. Koller K, Rudrapatna U, Chamberland M, Raven EP, Parker GD, Tax CMW, et al. MICRA: Microstructural image compilation with repeated acquisitions. Neuroimage. 2021 Jan 15;225:117406.

9. Jones DK, Alexander DC, Bowtell R, Cercignani M, Dell’Acqua F, McHugh DJ, et al. Microstructural imaging of the human brain with a “super-scanner”: 10 key advantages of ultra-strong gradients for diffusion MRI. Neuroimage. 2018 Nov 15;182:8–38.

10. Veraart J, Poot DHJ, Van Hecke W, Blockx I, Van der Linden A, Verhoye M, et al. More accurate estimation of diffusion tensor parameters using diffusion Kurtosis imaging. Magn Reson Med. 2011 Jan;65(1):138–45.

11. Jang I, Chun IY, Brosch JR, Bari S, Zou Y, Cummiskey BR, et al. Every hit matters: White matter diffusivity changes in high school football athletes are correlated with repetitive head acceleration event exposure. Neuroimage Clin. 2019 Jul 16;24:101930.

12. Palacios EM, Yuh EL, Mac Donald CL, Bourla I, Wren-Jarvis J, Sun X, et al. Diffusion Tensor Imaging Reveals Elevated Diffusivity of White Matter Microstructure that Is Independently Associated with Long-Term Outcome after Mild Traumatic Brain Injury: A TRACK-TBI Study. J Neurotrauma. 2022 Oct;39(19–20):1318–28.

13. Huang S, Huang C, Li M, Zhang H, Liu J. White matter abnormalities and cognitive deficit after mild traumatic brain injury: comparing DTI, DKI, and NODDI. Front Neurol. 2022 Mar 10;13:803066.

14. Liu Y, Lu L, Li F, Chen Y-C. Neuropathological mechanisms of mild traumatic brain injury: A perspective from multimodal magnetic resonance imaging. Front Neurosci. 2022 Jun 17;16:923662.

15. Jensen JH, Helpern JA, Ramani A, Lu H, Kaczynski K. Diffusional kurtosis imaging: the quantification of non-gaussian water diffusion by means of magnetic resonance imaging. Magnetic Resonance in Medicine: An Official Journal of the International Society for Magnetic Resonance in Medicine. 2005;53(6):1432–40.

16. Grossman EJ, Jensen JH, Babb JS, Chen Q, Tabesh A, Fieremans E, et al. Cognitive impairment in mild traumatic brain injury: a longitudinal diffusional kurtosis and perfusion imaging study. AJNR Am J Neuroradiol. 2013 May;34(5):951–7, S1.

17. Stenberg J, Skandsen T, Moen KG, Vik A, Eikenes L, Håberg AK. Diffusion Tensor and Kurtosis Imaging Findings the First Year following Mild Traumatic Brain Injury. J Neurotrauma. 2023 Mar;40(5–6):457–71.

18. Stokum JA, Sours C, Zhuo J, Kane R, Shanmuganathan K, Gullapalli RP. A longitudinal evaluation of diffusion kurtosis imaging in patients with mild traumatic brain injury. Brain Inj. 2015;29(1):47–57.

19. Funk JR, Duma SM, Manoogian SJ, Rowson S. Biomechanical risk estimates for mild traumatic brain injury. Annu Proc Assoc Adv Automot Med. 2007;51:343–61.

20. Rowson S, Duma SM, Beckwith JG, Chu JJ, Greenwald RM, Crisco JJ, et al. Rotational head kinematics in football impacts: an injury risk function for concussion. Ann Biomed Eng. 2012 Jan;40(1):1–13.

21. Greybe DG, Jones CM, Brown MR, Williams EMP. Comparison of head impact measurements via an instrumented mouthguard and an anthropometric testing device. Sports Eng. 2020 Dec;23(1):12.

22. Wu LC, Nangia V, Bui K, Hammoor B, Kurt M, Hernandez F, et al. In vivo evaluation of wearable head impact sensors. Ann Biomed Eng. 2016 Apr;44(4):1234–45.

23. Barnes-Wood M, McCloskey H, Connelly S, Gilchrist MD, Ni Annaidh A, Theobald P. Investigation of Head Kinematics and Brain Strain Response During Soccer Heading Using a Custom-Fit Instrumented Mouth guard. Currently Under Consideration with Annals of Biomedical Engineering. 2023;

24. Jones CM, Austin K, Augustus SN, Nicholas KJ, Yu X, Baker C, et al. An Instrumented Mouthguard for Real-Time Measurement of Head Kinematics under a Large Range of Sport Specific Accelerations. Sensors. 2023 Aug 10;23(16).

25. Smith SM. Fast robust automated brain extraction. Hum Brain Mapp. 2002 Nov;17(3):143–55.

26. Cordero-Grande L, Christiaens D, Hutter J, Price AN, Hajnal JV. Complex diffusion-weighted image estimation via matrix recovery under general noise models. Neuroimage. 2019 Oct 15;200:391–404.

27. Veraart J, Fieremans E, Novikov DS. Diffusion MRI noise mapping using random matrix theory. Magn Reson Med. 2016;76(5):1582–93.

28. Tournier J-D, Smith R, Raffelt D, Tabbara R, Dhollander T, Pietsch M, et al. MRtrix3: A fast, flexible and open software framework for medical image processing and visualisation. Neuroimage. 2019 Nov 15;202:116137.

29. Sairanen V, Leemans A, Tax CMW. Fast and accurate Slicewise OutLIer Detection (SOLID) with informed model estimation for diffusion MRI data. Neuroimage. 2018 Nov 1;181:331–46.

30. Andersson JLR, Skare S, Ashburner J. How to correct susceptibility distortions in spin-echo echo-planar images: application to diffusion tensor imaging. Neuroimage. 2003 Oct;20(2):870–88.

31. Smith SM, Jenkinson M, Woolrich MW, Beckmann CF, Behrens TEJ, Johansen-Berg H, et al. Advances in functional and structural MR image analysis and implementation as FSL. Neuroimage. 2004;23 Suppl 1:S208–19.

32. Andersson JLR, Sotiropoulos SN. An integrated approach to correction for off-resonance effects and subject movement in diffusion MR imaging. Neuroimage. 2016 Jan 15;125:1063–78.

33. Kellner E, Dhital B, Kiselev VG, Reisert M. Gibbs-ringing artifact removal based on local subvoxel-shifts. Magn Reson Med. 2016;76(5):1574–81.

34. Jeurissen B, Tournier J-D, Dhollander T, Connelly A, Sijbers J. Multi-tissue constrained spherical deconvolution for improved analysis of multi-shell diffusion MRI data. Neuroimage. 2014 Dec;103:411–26.

35. Ling J, Merideth F, Caprihan A, Pena A, Teshiba T, Mayer AR. Head injury or head motion? Assessment and quantification of motion artifacts in diffusion tensor imaging studies. Hum Brain Mapp. 2012 Jan;33(1):50–62.

36. Neto Henriques R, Correia MM, Nunes RG, Ferreira HA. Exploring the 3D geometry of the diffusion kurtosis tensor--impact on the development of robust tractography procedures and novel biomarkers. Neuroimage. 2015 May 1;111:85–99.

37. Wasserthal J, Neher P, Maier-Hein KH. TractSeg - Fast and accurate white matter tract segmentation. Neuroimage. 2018 Dec;183:239–53.

38. Chandio BQ, Risacher SL, Pestilli F, Bullock D, Yeh F-C, Koudoro S, et al. Bundle analytics, a computational framework for investigating the shapes and profiles of brain pathways across populations. Sci Rep. 2020 Oct 13;10(1):17149.

39. Zeng W. lmerPerm: Perform Permutation Test on General Linear and Mixed Linear Regression. CRAN; 2023.

40. Virtanen P, Gommers R, Oliphant TE, Haberland M, Reddy T, Cournapeau D, et al. SciPy 1.0: fundamental algorithms for scientific computing in Python. Nat Methods. 2020 Mar;17(3):261–72.

41. Hunter JD. Matplotlib: A 2D Graphics Environment. Comput Sci Eng. 2007;9(3):90–5.

42. Gommers R, Virtanen P, Burovski E, Weckesser W, Oliphant TE, Haberland M, et al. scipy/scipy: SciPy 1.9. 0. Zenodo. 2022;

43. Kenny RA, Mayo CD, Kennedy S, Varga AA, Stuart-Hill L, Garcia-Barrera MA, et al. A pilot study of diffusion tensor imaging metrics and cognitive performance pre and post repetitive, intentional sub-concussive heading in soccer practice. Journal of Concussion. 2019 Jan;3:205970021988550.

44. Brett BL, Bobholz SA, España LY, Huber DL, Mayer AR, Harezlak J, et al. Cumulative Effects of Prior Concussion and Primary Sport Participation on Brain Morphometry in Collegiate Athletes: A Study From the NCAA-DoD CARE Consortium. Front Neurol. 2020 Jul 28;11:673.

45. Churchill NW, Hutchison MG, Di Battista AP, Graham SJ, Schweizer TA. Structural, Functional, and Metabolic Brain Markers Differentiate Collision versus Contact and Non-Contact Athletes. Front Neurol. 2017 Aug 22;8:390.

46. Goubran M, Mills BD, Georgiadis M, Karimpoor M, Mouchawar N, Sami S, et al. Microstructural alterations in tract development in college football and volleyball players: A longitudinal diffusion MRI study. Neurology. 2023 Aug 29;101(9):e953–65.

47. Kwiatkowski A, Weidler C, Habel U, Coverdale NS, Hirad AA, Manning KY, et al. Uncovering the hidden effects of repetitive subconcussive head impact exposure: A mega-analytic approach characterizing seasonal brain microstructural changes in contact and collision sports athletes. Hum Brain Mapp. 2024 Aug 15;45(12):e26811.

48. Mustafi SM, Harezlak J, Koch KM, Nencka AS, Meier TB, West JD, et al. Acute White-Matter Abnormalities in Sports-Related Concussion: A Diffusion Tensor Imaging Study from the NCAA-DoD CARE Consortium. J Neurotrauma. 2018 Nov 15;35(22):2653–64.

49. Chung S, Chen J, Li T, Wang Y, Lui YW. Investigating Brain White Matter in Football Players with and without Concussion Using a Biophysical Model from Multishell Diffusion MRI. AJNR Am J Neuroradiol. 2022 Jun;43(6):823–8.

50. Wang ML, Yu MM, Yang DX, Liu YL, Wei XE, Li WB. Longitudinal microstructural changes in traumatic brain injury in rats: A diffusional kurtosis imaging, histology, and behavior study. AJNR Am J Neuroradiol. 2018 Sep;39(9):1650–6.

51. Karlsen RH, Einarsen C, Moe HK, Håberg AK, Vik A, Skandsen T, et al. Diffusion kurtosis imaging in mild traumatic brain injury and postconcussional syndrome. J Neurosci Res. 2019 May;97(5):568–81.

52. Tayebi M, Holdsworth SJ, Champagne AA, Cook DJ, Nielsen P, Lee T-R, et al. The role of diffusion tensor imaging in characterizing injury patterns on athletes with concussion and subconcussive injury: a systematic review. Brain Inj. 2021 May 12;35(6):621–44.

53. Wang F, Wang Z, Hu L, Xu H, Yu C, Li F. Evaluation of Head Injury Criteria for Injury Prediction Effectiveness: Computational Reconstruction of Real-World Vulnerable Road User Impact Accidents. Front Bioeng Biotechnol. 2021 Jun 29;9:677982.

54. Alshareef A, Giudice JS, Forman J, Shedd DF, Reynier KA, Wu T, et al. Biomechanics of the Human Brain during Dynamic Rotation of the Head. J Neurotrauma. 2020 Jul 1;37(13):1546–55.

55. Abbasi Ghiri A, Seidi M, Wallace J, Cheever K, Memar M. Exploring Sex-Based Variations in Head Kinematics During Soccer Heading. Annals of Biomedical Engineering. 2025;1–17.

56. Kern J, Hermsdörfer J, Gulde P. Factors Influencing Wearable-Derived Head Impact Kinematics in Soccer Heading. German Journal of Sports Medicine/Deutsche Zeitschrift fur Sportmedizin. 2024;75(3).

57. Bradbery E, Cairns R, Peek K. The relationship between header type and head acceleration during heading in male youth football players. Physical Therapy in Sport. 2024;

58. Kimpara H, Iwamoto M. Mild traumatic brain injury predictors based on angular accelerations during impacts. Ann Biomed Eng. 2012 Jan;40(1):114–26.

