## Supplementary Figure for "LONGITUDINAL CHANGES IN WHITE MATTER MICROSTRUCTURAL STATUS FOLLOWING QUANTIFIED HEAD-BALL IMPACTS IN SOCCER: A PRELIMINARY, PROSPECTIVE STUDY"

**SUPPLEMENTARY MATERIAL**

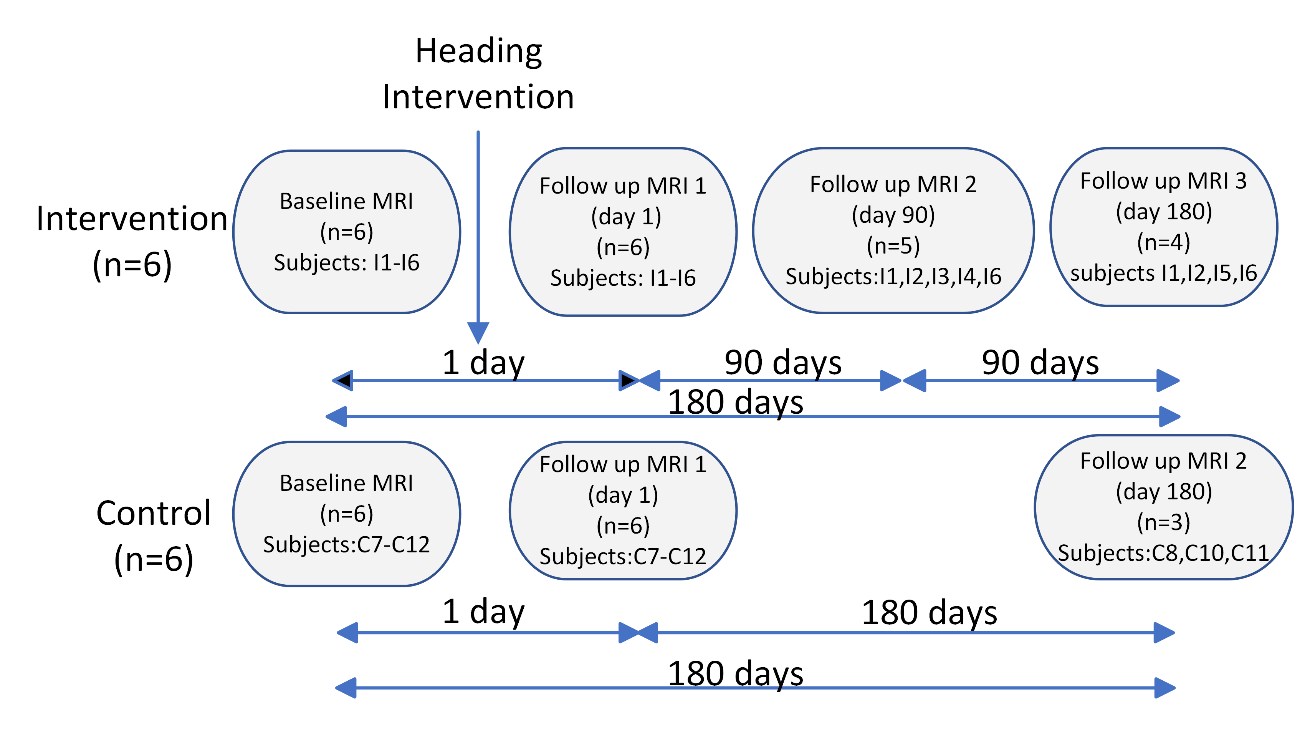

**Supplementary Figure 1**

Pictorial explanation of participant engagement and retention, relative to scanning sessions.

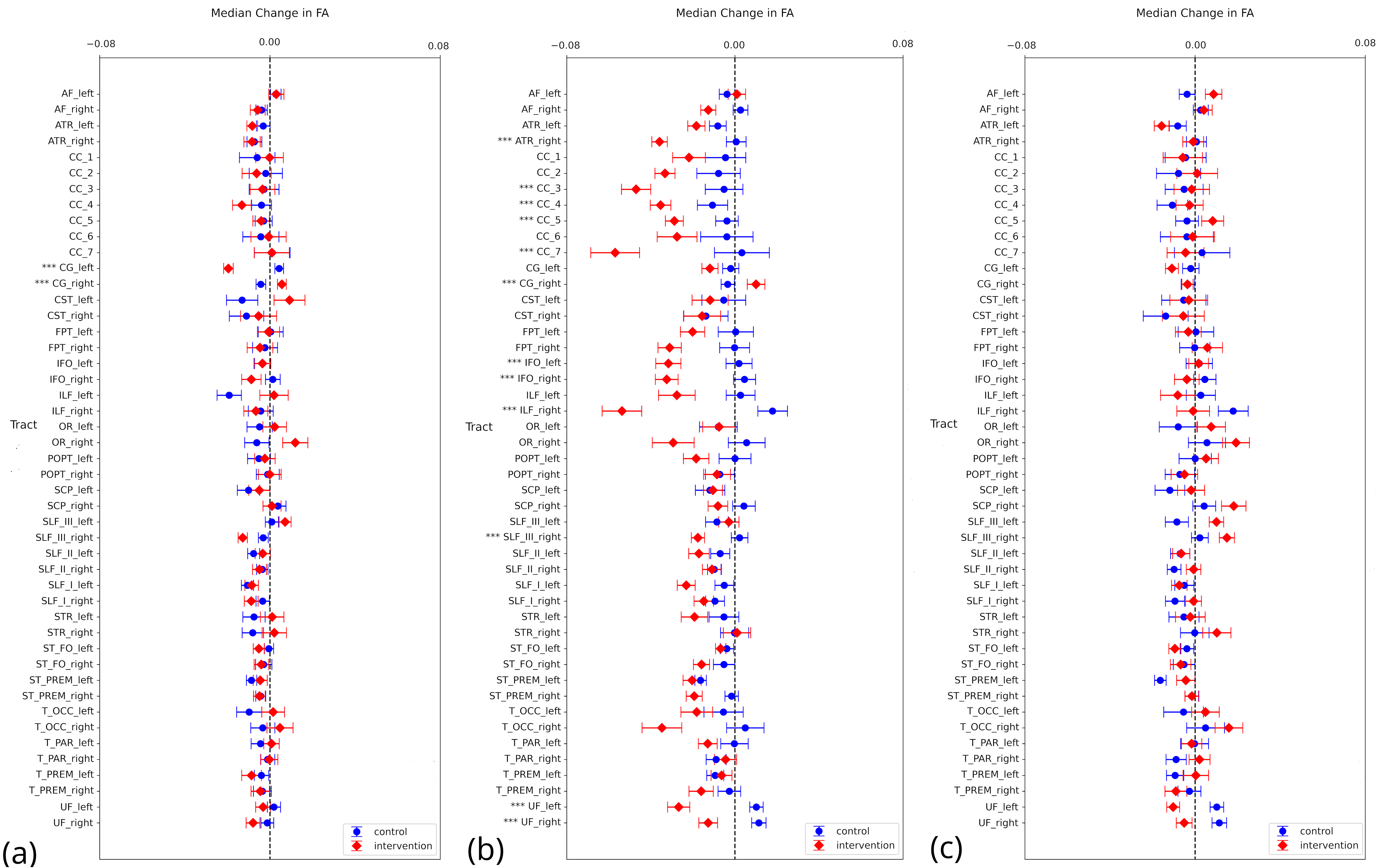

**Supplementary Figure 2**

Variation in FA between day 1 (a), day 90 (b) and day (180), relative to day 0 measures. Those tracts that have a statistically significant difference (p < 0.05) between the Control and Intervention metrics are signified by ***.

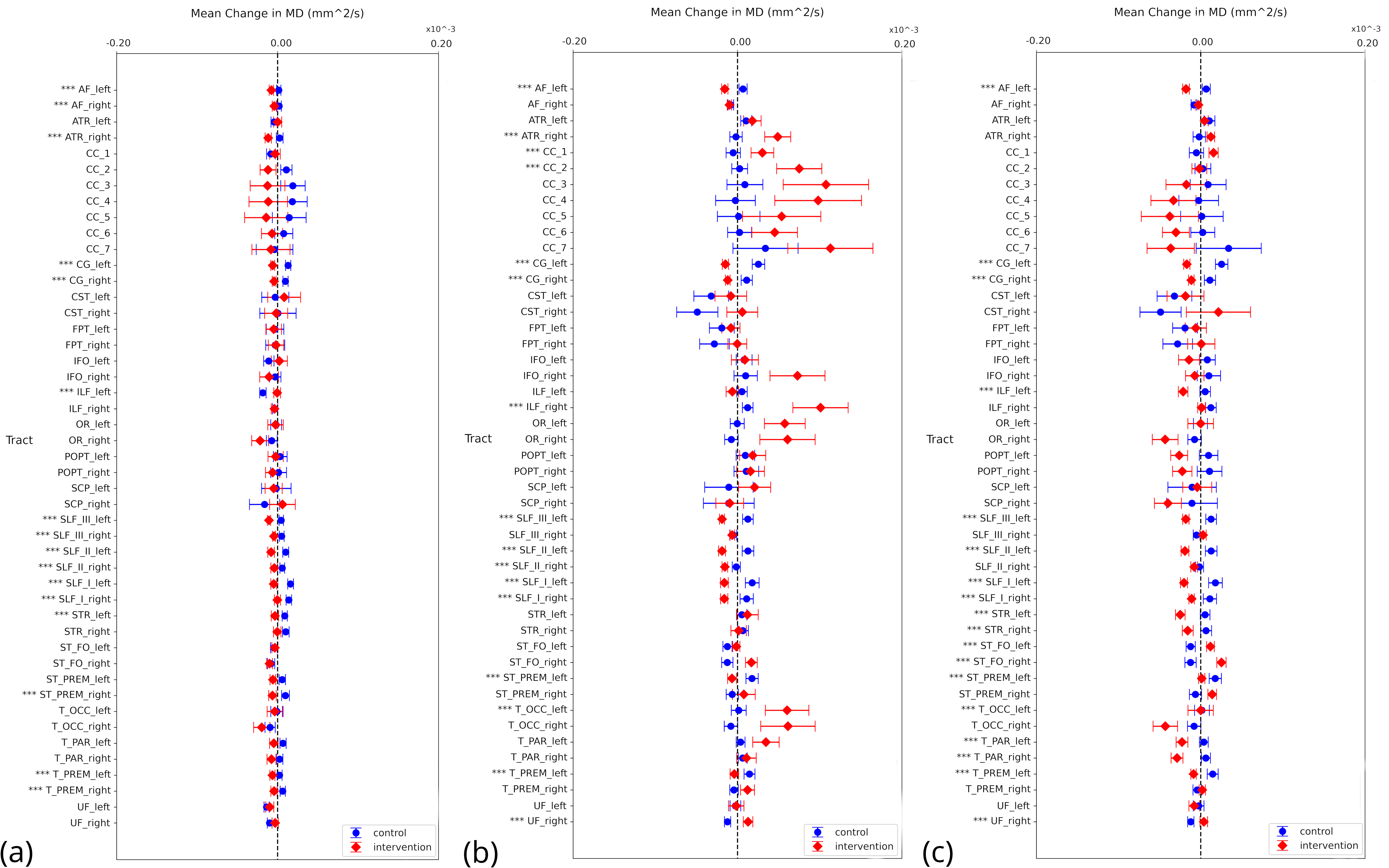

**Supplementary Figure 3:** Variation in MD between day 1 (a), day 90 (b) and day (180), relative to day 0 measures. Those tracts that have a statistically significant difference (p < 0.05) between the Control and Intervention metrics are signified by ***.

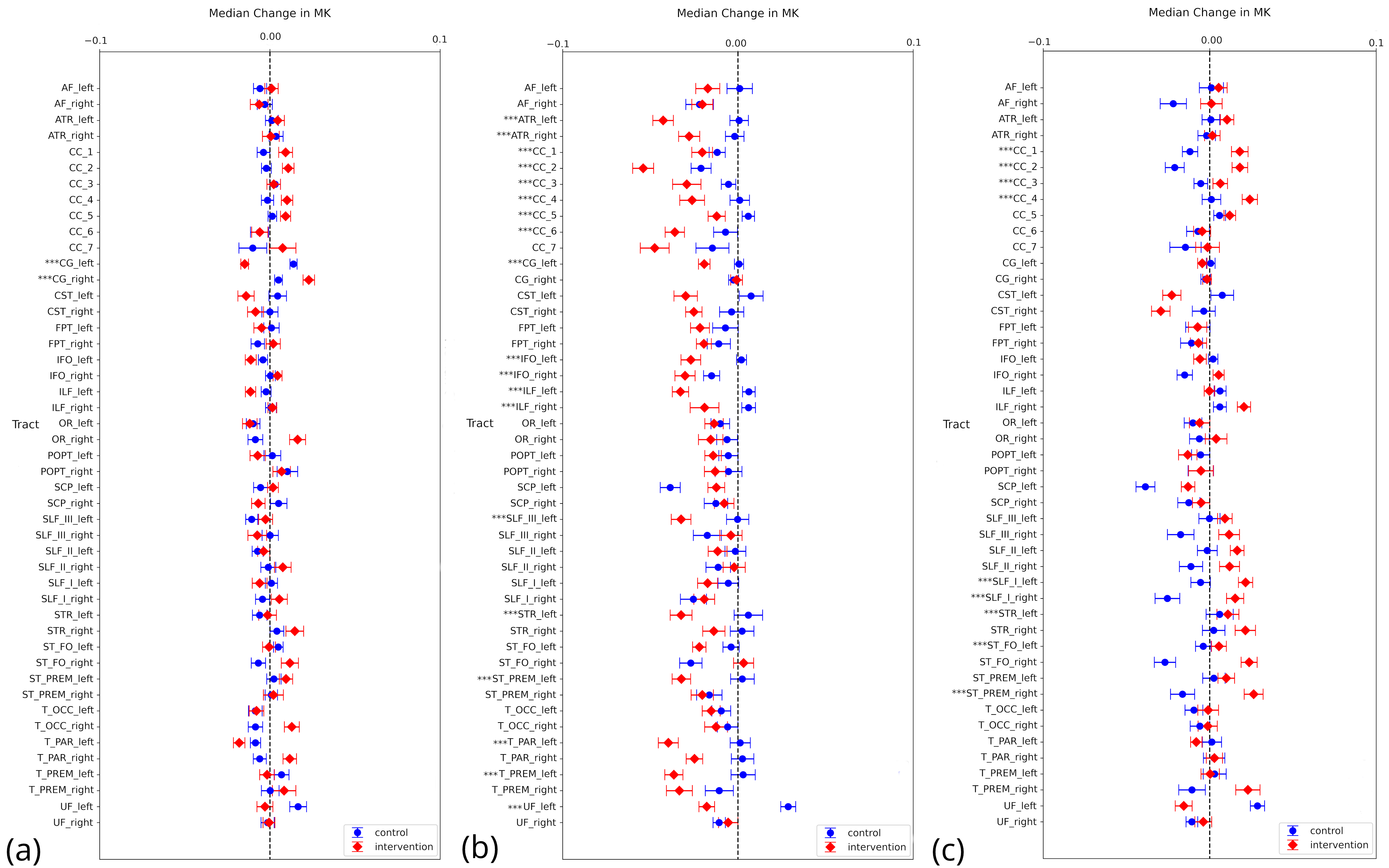

**Supplementary Figure 4:** Variation in MK between day 1 (a), day 90 (b) and day (180), relative to day 0 measures. Those tracts that have a statistically significant difference (p < 0.05) between the Control and Intervention metrics are signified by ***.

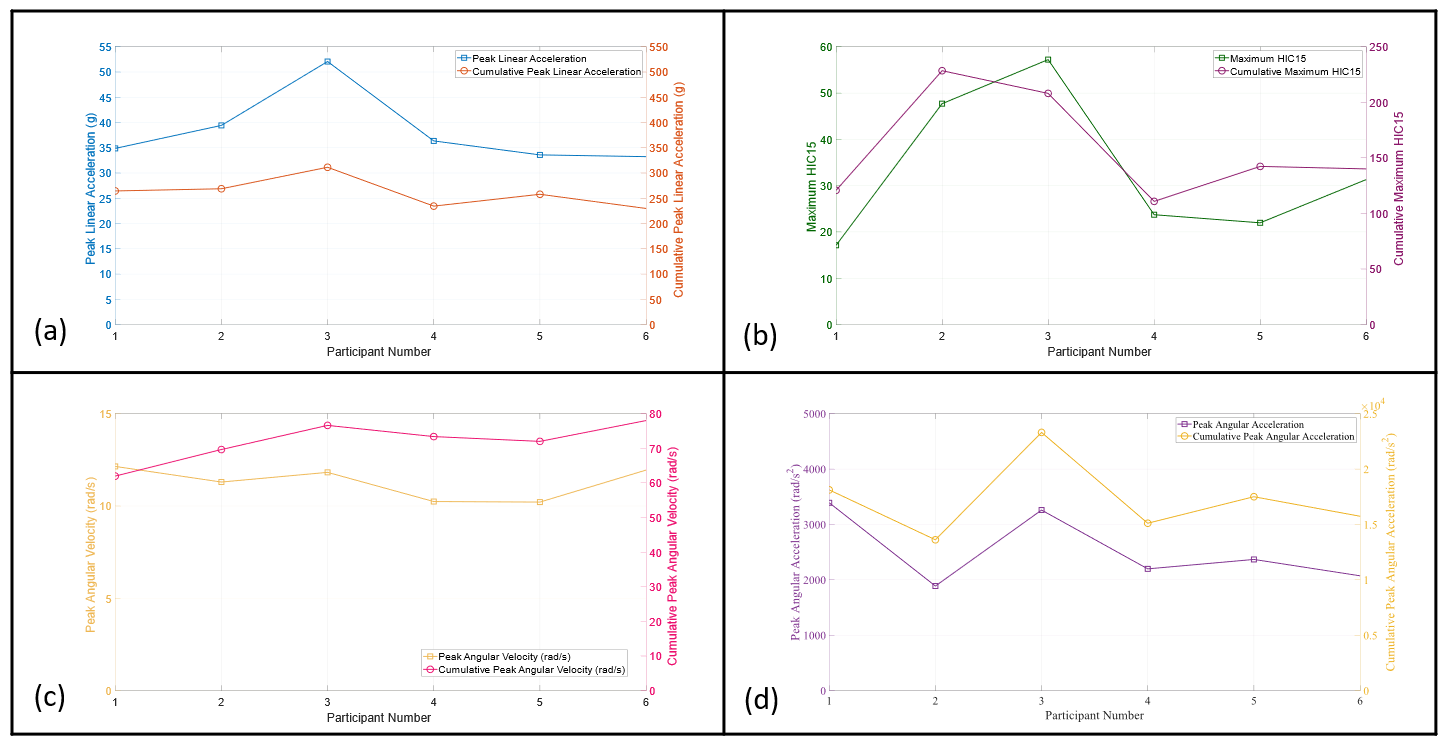

**Supplementary Figure 5**

The peak kinematics reported for each Intervention participant. (a) Peak linear acceleration (blue) and cumulative PLA (orange); (b) Peak and cumulative head injury criterion scores, calculated over a 15 ms window about the peak acceleration; (c) Peak angular velocity and cumulative PAV; (d) Peak angular acceleration and cumulative PAA.

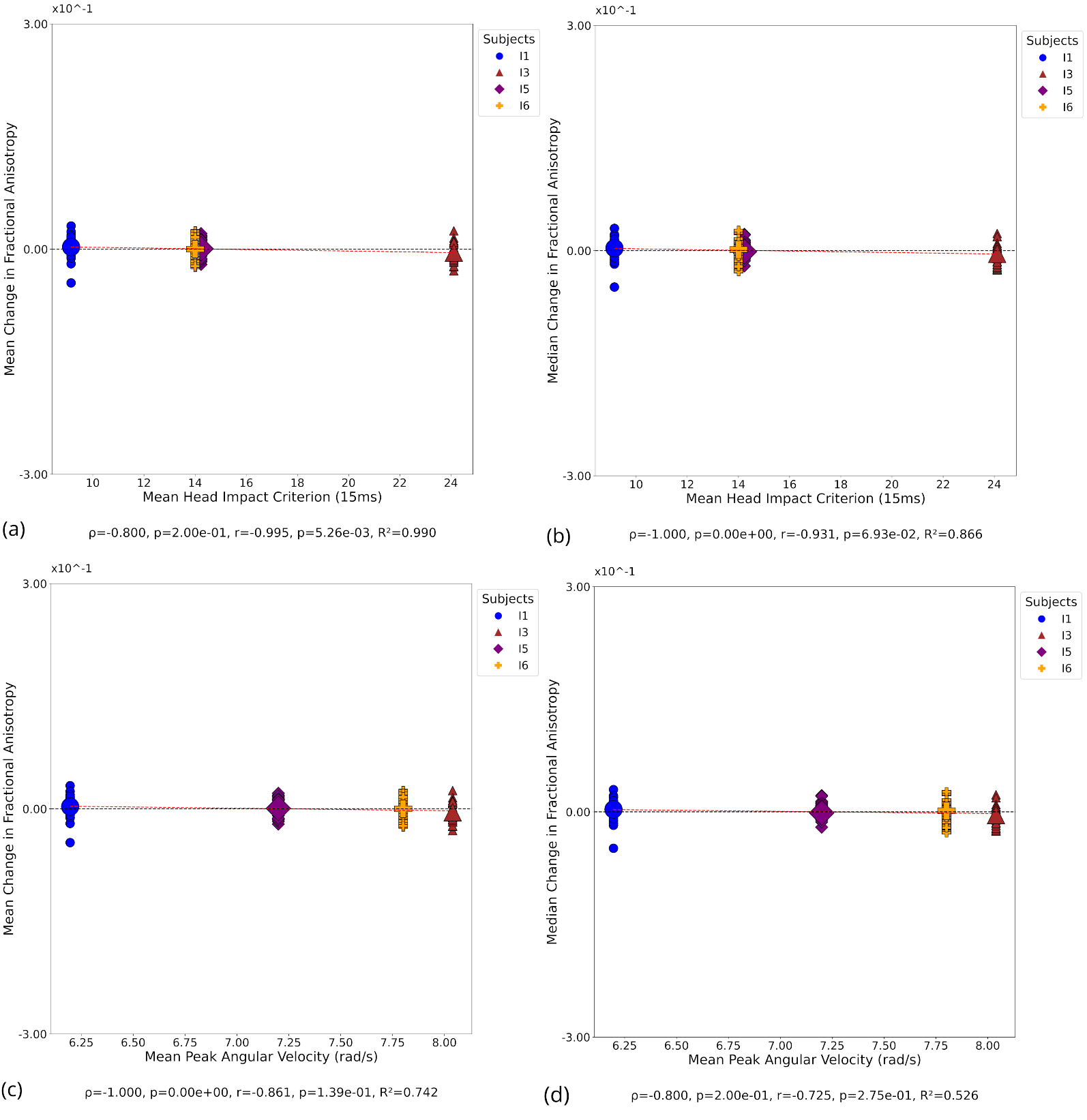

**Supplementary Figure 6**

Correlation with FA at day 180. Small symbols indicate change in FA in individual white matter tracts at the participant level; large symbols indicate overall change in FA for each participant**.** Pearson's correlation (r), Spearman's correlation (ρ), and coefficient of determination (R²).  **(a)** Mean change in FA versus mean HIC. **(b)** Median change in FA versus mean HIC. **(c)** Mean change in FA versus mean PAV. **(d)** Median change in FA versus mean PAV.

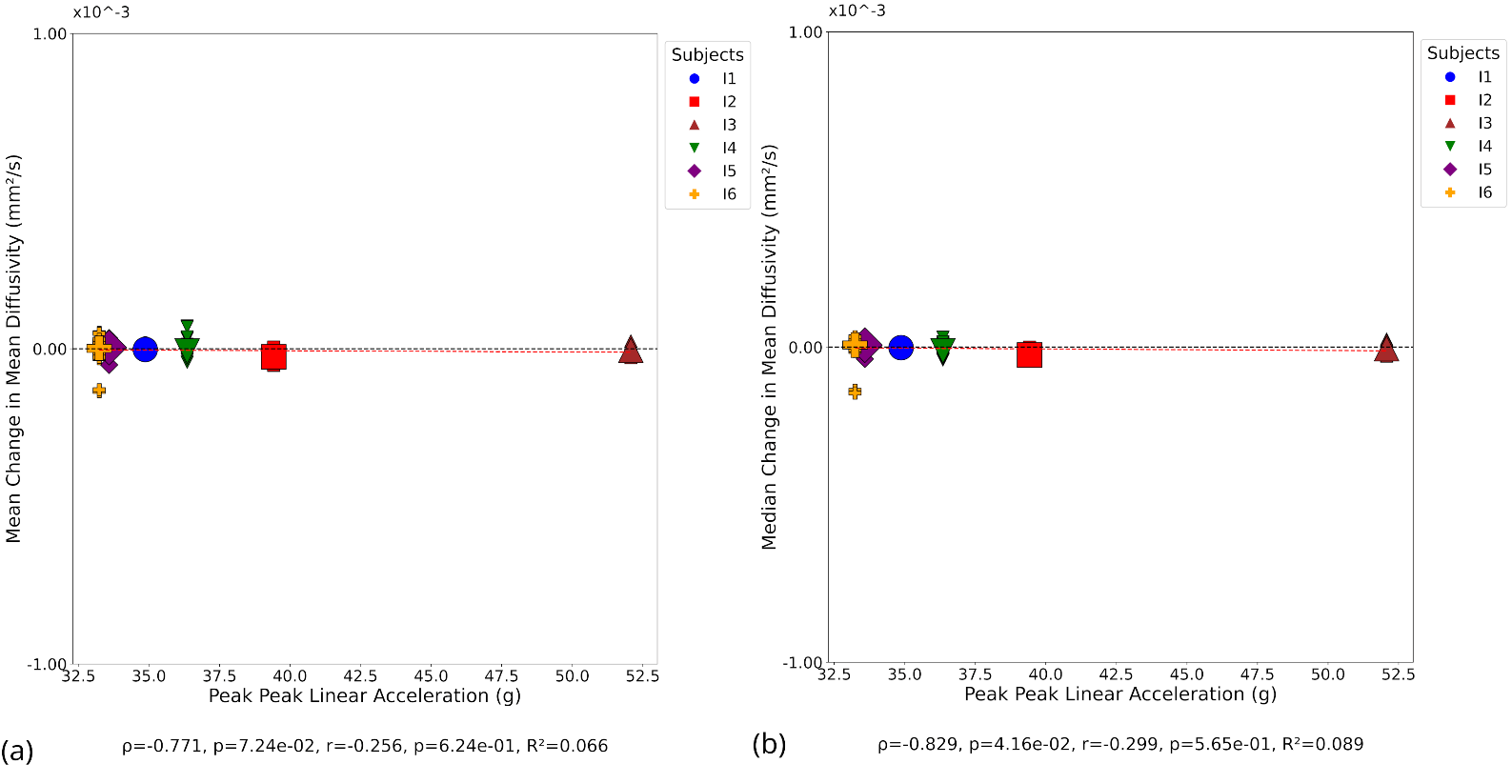

**Supplementary Figure 7**

Correlation with MD at day 1. Small symbols indicate change in MD in individual white matter tracts at the participant level; large symbols indicate overall change in MD for each participant**.** Pearson's correlation (r), Spearman's correlation (ρ), and coefficient of determination (R²).  **(a)** Mean change in MD versus mean HIC. **(b)** Median change in MD versus mean HIC. **(c)** Mean change in MD versus mean PAV. **(d)** Median change in MD versus mean PAV.

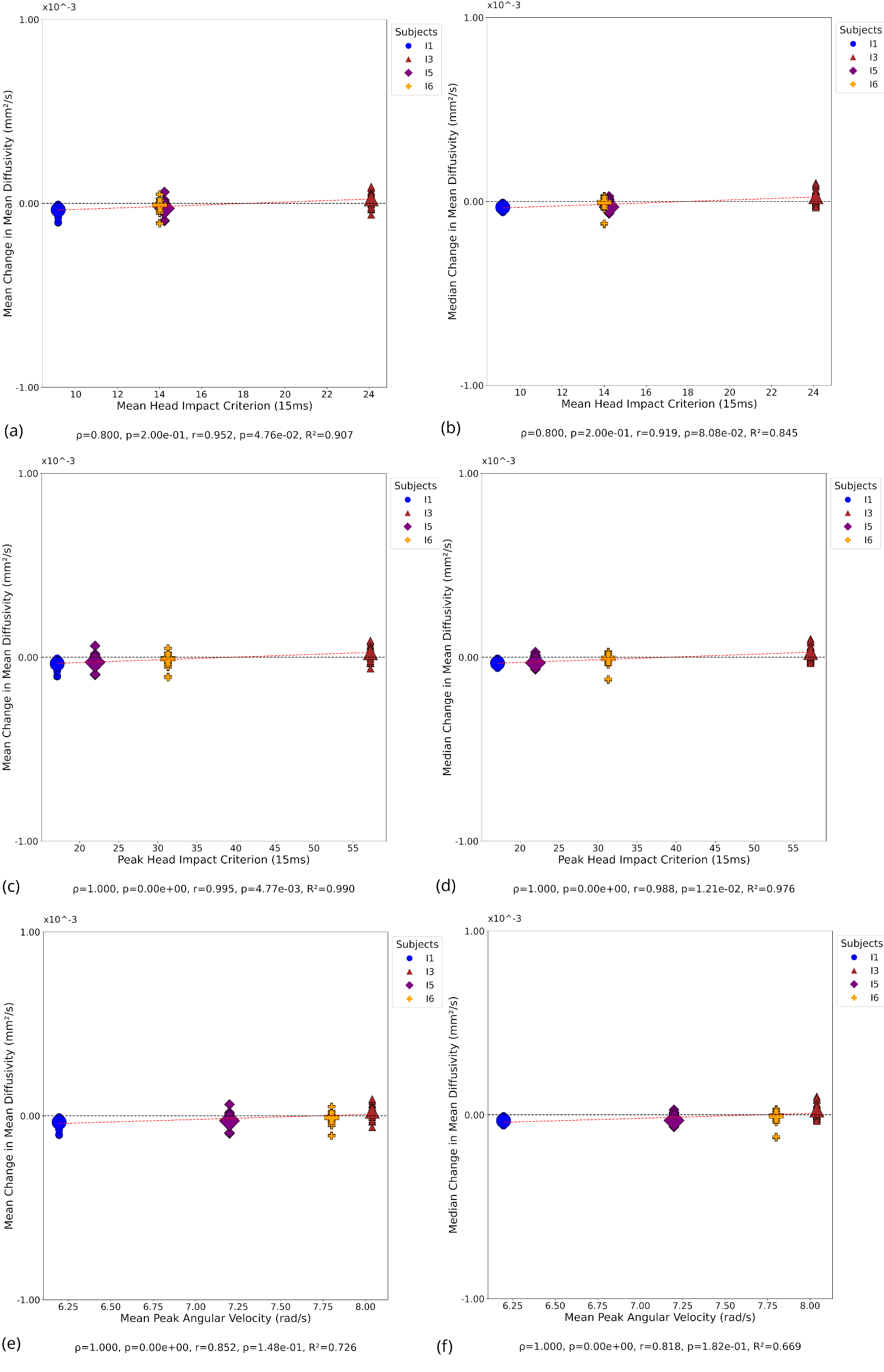

**Supplementary Figure 8:**

Correlation with MD at day 180. Small symbols indicate change in MD in individual white matter tracts at the participant level; large symbols indicate overall change in MD for each participant**.** Pearson's correlation (r), Spearman's correlation (ρ), and coefficient of determination (R²).  **(a)** Mean change in MD versus mean HIC. **(b)** Median change in MD versus mean HIC. **(c)** Mean change in MD versus peak HIC. **(d)** Median change in MD versus peak HIC. **(e)** Mean change in MD versus mean PAV. **(f)** Median change in MD versus mean PAV.

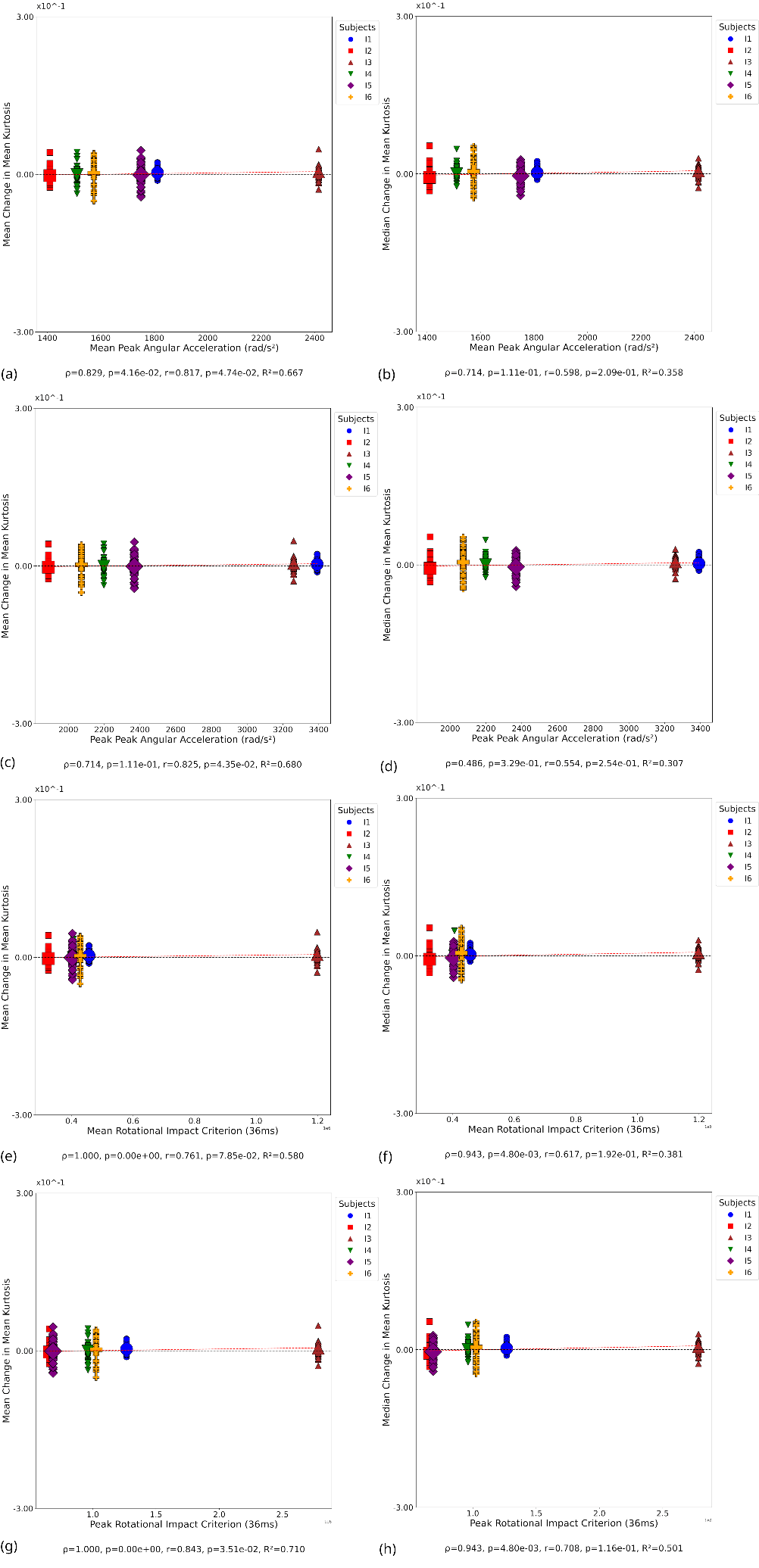

**Supplementary Figure 9**

Correlation with MK at day 180. Small symbols indicate change in MD in individual white matter tracts at the participant level; large symbols indicate overall change in MK for each participant**.** Pearson's correlation (r), Spearman's correlation (ρ), and coefficient of determination (R²).  **(a)** Mean change in MK versus mean PAA. **(b)** Median change in MK versus mean PAA. **(c)** Mean change in MK versus peak PAA. **(d)** Median change in MK versus peak PAA.
