## Supplementary Table for "LONGITUDINAL CHANGES IN WHITE MATTER MICROSTRUCTURAL STATUS FOLLOWING QUANTIFIED HEAD-BALL IMPACTS IN SOCCER: A PRELIMINARY, PROSPECTIVE STUDY"

**Supplementary Table 1:** A list of the TractSeg labels and associated anatomical tract names.

| | |
|---|---|
| AF_left | Arcuate fascicle left |
| AF_right | Arcuate fascicle right |
| ATR_left | Anterior Thalamic Radiation |
| ATR_right | Anterior Thalamic Radiation right |
| CC_1 | Corpus Callosum Rostrum |
| CC2 | Corpus Callosum Genu |
| CC3 (Premotor) | Corpus Callosum Rostral body (Premotor) |
| CC4 | Corpus Callosum Anterior midbody (Primary Motor) |
| CC5 | Corpus Callosum Posterior midbody (Primary Somatosensory) |
| CC6 | Corpus Callosum Isthmus |
| CC7 | Corpus Callosum Splenium |
| CG_left | Cingulum left |
| CG_right | Cingulum right |
| CST_left | Corticospinal tract left |
| CST_right | Corticospinal tract right |
| FPT_left | Fronto-pontine tract left |
| FPT_right | Fronto-pontine tract right |
| ICP_left | Inferior cerebellar peduncle left |
| ICP_right | Inferior cerebellar peduncle right |
| IFO_left | Inferior occipito-frontal fascicle left |
| IFO_right | Inferior occipito-frontal fascicle right |
| ILF_left | Inferior longitudinal fascicle left |
| ILF_right | Inferior longitudinal fascicle right |
| MCP | Middle cerebellar peduncle |
| OR_left | Optic radiation left |
| OR_right | Optic radiation right |
| POPT_left | Parieto-occipital pontine left |
| POPT_right | Parieto-occipital pontine right |
| SCP_left | Superior cerebellar peduncle left |
| SCP_right | Superior cerebellar peduncle right |
| SLF_I_left | Superior longitudinal fascicle I left |
| SLF_I_right | Superior longitudinal fascicle I right |
| SLF_II_left | Superior longitudinal fascicle II left |
| SLF_II_right | Superior longitudinal fascicle II right |
| SLF_III_left | Superior longitudinal fascicle III left |
| SLF_III_right | Superior longitudinal fascicle III right |
| STR_left | Superior Thalamic Radiation left |
| STR_right | Superior Thalamic Radiation right |
| UF_left | Uncinate fascicle left |
| UF_right | Uncinate fascicle right |
| T_PREM_left | Thalamo-premotor left |
| T_PREM_right | Thalamo-premotor right |
| T_PAR_left | Thalamo-parietal left |
| T_PAR_right | Thalamo-parietal right |
| T_OCC_left | Thalamo-occipital left |
| T_OCC_right | Thalamo-occipital right |
| ST_FO_left | Striato-fronto-orbital left |
| ST_FO_right | Striato-fronto-orbital right |
| ST_PREM_left | Striato-premotor left |
| ST_PREM_right | Striato-premotor right |
| ST_PAR_left | Striato-parietal left |
| ST_PAR_right | Striato-parietal right |
| ST_OCC_left | Striato-occipital left |
| ST_OCC_right | Striato-occipital right |
